# Identification and characterization of novel proteins from Arizona bark scorpion venom that inhibit Nav1.8, a voltage-gated sodium channel regulator of pain signaling

**DOI:** 10.1101/2021.06.24.449822

**Authors:** Tarek Mohamed Abd El-Aziz, Yucheng Xiao, Jake Kline, Harold Gridley, Alyse Heaston, Klaus D. Linse, Micaiah J. Ward, Darin R. Rokyta, James D. Stockand, Theodore R. Cummins, Luca Fornelli, Ashlee H. Rowe

## Abstract

The voltage-gated sodium channel Nav1.8 is linked to neuropathic and inflammatory pain, highlighting the potential to serve as a drug target. However, the biophysical mechanisms that regulate Nav1.8 activation and inactivation gating are not completely understood. Progress has been hindered by a lack of biochemical tools for examining Nav1.8 gating mechanisms. Arizona bark scorpion (*Centruroides sculpturatus*) venom proteins inhibit Nav1.8 and block pain in grasshopper mice (*Onychomys torridus*). These proteins provide tools for examining Nav1.8 structure-activity relationships. To identify proteins that inhibit Nav1.8 activity, venom samples were fractioned using liquid chromatography (reversed phase and ion exchange). A recombinant Nav1.8 clone expressed in ND7/23 cells was used to identify subfractions that inhibited Nav1.8 Na+ current. Mass spectrometry-based bottom-up proteomic analyses identified unique peptides from inhibitory subfractions. A search of the peptides against the AZ bark scorpion venom gland transcriptome revealed four novel proteins between 40 and 60% conserved with venom proteins from scorpions in four genera (*Centruroides*, *Parabuthus*, *Androctonus*, and *Tityus*). Ranging from 63 to 82 amino acids, each primary structure includes 8 cysteines and a “CXCE” motif where X = an aromatic residue (tryptophan, tyrosine or phenylalanine). Electrophysiology data demonstrated that the inhibitory effects of bioactive subfractions can be removed by hyperpolarizing the channels, suggesting that proteins may function as gating modifiers as opposed to pore blockers.

## 1. Introduction

Primary sensory nociceptive neurons, whose cell bodies reside in the dorsal root ganglia (DRG) of the body and limbs, or trigeminal ganglia (TG) of the head and face, transmit pain signals to the central nervous system (CNS) [1–4]. Tissue damage due to injury, aging or disease causes biochemical changes in nociceptive neurons that activate the voltage-gated sodium channel (VGSC) Nav1.7, which then recruits Nav1.8 [1,5–8]. Activation of Nav1.8 generates the majority of Na^+^ current underlying the action potentials that carry pain signals to the brain; inactivation terminates the action-potential driven pain signals [5–8]. Chronic pain results from prolonged activation or failed inactivation [1,9]. Nav1.8 activity is linked to mechanical, neuropathic and inflammatory pain, highlighting the potential for Nav1.8 to serve as a drug target [7,8,10–18]. However, the biophysical mechanisms that regulate Nav1.8 gating are not completely understood, particularly the mechanisms that regulate Nav1.8 inactivation [9].

The function of a VGSC is to open briefly, allowing an influx of Na^+^ ions to depolarize cell membranes, generating the action potentials that underlie neuronal signaling and muscle contraction [19–21]. VGSC alpha subunits have four domains, DI - DIV (**Figure 1**). Each domain has 6 membrane-spanning helices (S1 – S6), and re-entrant loops that connect S5 and S6 (Pore). The domains are organized such that the re-entrant loops face each other to form an ion-permeating pore with an activation gate. When a cell is at rest, the activation gate is closed. The S4 segments of each domain have alternating positively charged amino acids that function as voltage sensors. Biochemical changes in damaged tissues depolarize membranes, imposing electrostatic forces on these positive charges. The voltage sensors move outward and open the channel (activated) [20,22]. The outward movements of S4 in DI – DIII initiate channel activation (opening); the S4 in the fourth domain detects channel opening and initiates fast inactivation. During fast inactivation, part of the DIII – DIV intracellular loop forms a “hinged lid” inactivation gate that moves into the mouth of the channel to block the pore. A second type of inactivation, slow inactivation, occurs when the pore loops change conformation to block the flow of ions [23]. Failed inactivation prolongs the action potentials that carry pain signals to the brain.

**Figure 1.**
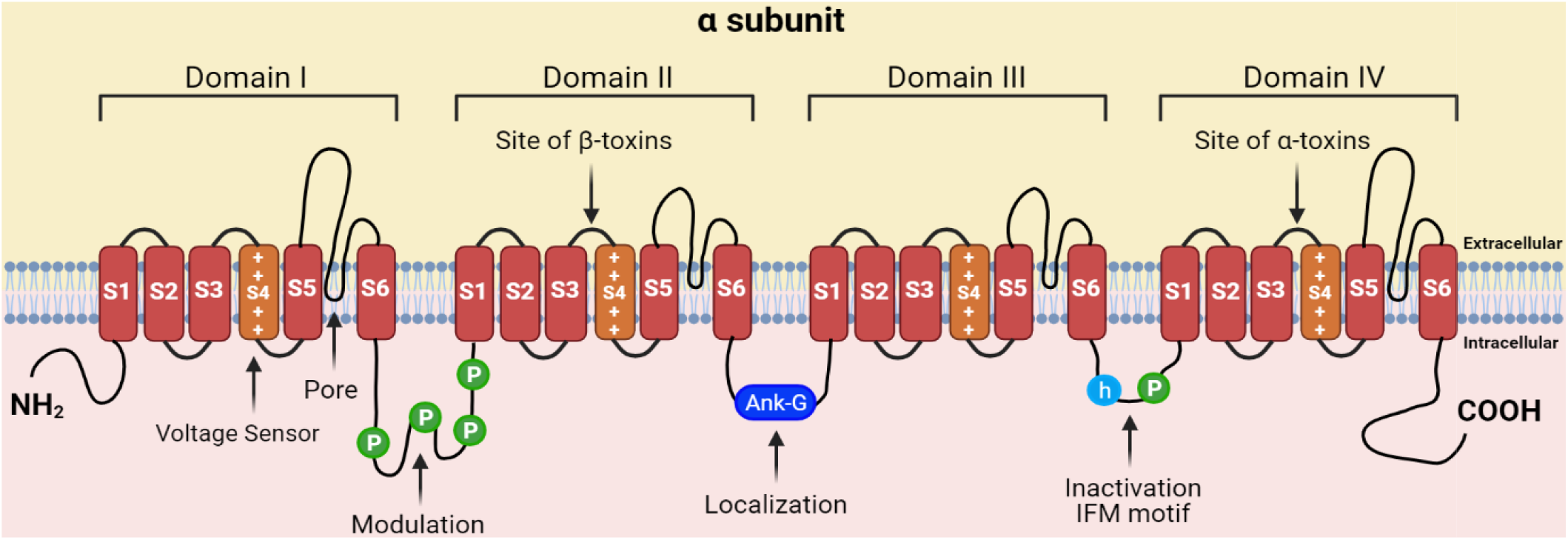
Structure of a voltage-gated sodium channel (VGSC). The channel consists of four repeating domains (DI-DIV), each having six transmembrane helices (S1-S6). Helix 4 (S4) in each domain serves as the voltage sensor. The S5-S6 re-entrant loops from the four domains join to form the pore. Re-entrant loops connecting S5 to S6 line the pore and form the Na+ selectivity filter and activation gate. When the cell is at rest, the activation gate is closed. When the channel is activated, the gate is open (the ball-and-chain inactivation gate is disengaged). During fast inactivation, the “hinged-lid” gate occludes the intracellular side of the pore. During slow inactivation, the pore loops change conformation to block the pore. Created with BioRender.com

Nav1.8 generates the action potentials that carry pain signals to the brain. Characterization of mechanisms that regulate Nav1.8 inactivation gating would advance efforts to engineer drugs that inhibit Nav1.8 activity and block the transmission of pain signals to the brain. Animal venoms are a rich source of proteins that bind and modulate nociceptive neurons [24–30]. Venoms from diverse animal taxa evolved to effectively subdue prey and deter predators. The venoms are composed of pharmacologically active cocktails containing salts, enzymatic and non-enzymatic proteins, and small molecules [31]. The development of new drugs is one of the most challenging activities in the pharmaceutical industry. Since the first animal venom-derived drug, Captopril, there are 10 other approved venom-derived drugs marketed commercially [31–33]. Five drugs were identified from snake venoms including enalapril, tirofiban, eptifibatide, batroxobin, and cobratide [31,34]. Exenatide and lixisenatide are derivatives isolated from lizards. Bivalirudin and desirudin are derivatives from leeches. Ziconotide, a non-opioid analgesic drug, is a synthetic version of ω-conotoxin MVIIA obtained from the venom of *Conus magus* [35]. Several other animal venom components are now involved in various clinical trials as new therapeutic molecules for medical purposes [32,36].

Proteins that modulate VGSC activity can serve as templates for structure-guided engineering of drugs that block pain [37–40]. Venom proteins bind the extracellular linkers adjacent to the voltage sensors that control channel activation and inactivation. Beta sodium toxin (*β*-NaTx) proteins trap the DII voltage sensor in the outward position, modifying channel activation. Alpha sodium toxin (α-NaTx) proteins trap the DIV voltage sensor in the inward position, preventing the fast inactivation hinged lid from docking with the pore. Because these proteins modify activation and fast inactivation gating, they are used as tools to examine VGSC structure-activity relationships [20,41–43]. For example, computational modeling of scorpion venom proteins bound to the DII voltage sensor were used to map voltage sensor structure and regulation of activation gating in Nav1.2, a brain VGSC [44–46]. Cryo-electron microscopy (cryo-EM) studies using scorpion venom proteins bound to Nav1.7, a channel responsible for spontaneous pain disorders, revealed the structural basis of fast inactivation [47]. Venom proteins that target Nav1.7 have also provided templates for structure-guided mutagenesis to engineer new pain drugs. For example, mutagenesis of a spider venom protein changed the native protein from a Nav1.7 channel activator to a channel inhibitor [39]. While Nav1.8 activity is linked to mechanical, neuropathic and inflammatory pain, highlighting the potential for Nav1.8 to serve as an alternative drug target to Nav1.7 [7,8,10–18], the biophysical mechanisms that regulate Nav1.8 gating are not completely understood, particularly inactivation. Progress has been hindered by a lack of venom proteins that modify Nav1.8 gating. In an effort to use gating modifier toxins to study Nav1.8 gating properties, Gilchrist and Bosmans exchanged Nav1.8 voltage sensors with Nav1.2, a toxin-sensitive channel [48]. Arizona (AZ) bark scorpion (*Centruroides sculpturatus*) venom proteins inhibit Nav1.8 activity and block pain in natural populations of predatory mice (Grasshopper mice, *Onychomys torridus*) that feed on scorpions [49]. Bark scorpions produce venom proteins that reversibly bind VGSCs [50–56]. Grasshopper mice show little response when either stung by scorpions or injected with venom [49,57,58]. Electrophysiological data showed that venom inhibited Nav1.8 currents and blocked action potentials in nociceptive neurons from mice [49]. Moreover, injections of venom into the hind paw of mice blocked pain-related behavior (paw licking) in response to subsequent injections of formalin, a chemical that induces pain. Thus, grasshopper mice provide a system for characterizing proteins that inhibit Nav1.8 activity and block pain signal transmission.

Our goal is to determine the molecular mechanisms underlying venom-mediated inhibition of grasshopper mice Nav1.8. AZ bark scorpion venom comprises multiple proteins. In order to isolate and identify proteins that inhibit grasshopper mice Nav1.8 activity, we used complementary methods including reversed phase (RPLC) and ion exchange (IEC) liquid chromatography, a recombinant Nav1.8 bioactivity assay, and proteomic analyses. Here, we report the validated mass and primary structure of four candidate proteins isolated from venom subfractions that inhibit grasshopper mice Nav1.8 Na^+^ current. In addition, we report that the Nav1.8-inhibiting effects of venom are voltage-dependent and can be removed by hyperpolarizing the channels. Similar to the electrophysiological properties of venom, the Nav1.8-inhibiting effects of subfractions isolated from venom are voltage dependent and can be removed by hyperpolarizing the channels. These findings are significant because few toxin proteins have been identified that modify Nav1.8 activity, particularly inhibition of activity [59]. Moreover, these findings provide a clue to the biophysical mechanism by which bark scorpion venom inhibits grasshopper mice Nav1.8. Voltage dependent effects suggest that venom proteins do not function as pore blockers and instead act as gating modifiers. These proteins potentially provide new tools for investigating the biophysical mechanisms underlying Nav1.8 activity, the first step toward engineering proteins that inhibit Nav1.8 activity in humans to block pain.

## 2. Results

### 2.1. RPLC fractionation of AZ bark scorpion venom

Reversed phase liquid chromatography (RPLC) was used to separate AZ bark scorpion venom into different fractions. Lyophilized AZ bark scorpion venom samples were hydrated in 0.1% trifluoroacetic acid in water and fractionated by reversed phase liquid chromatography using a C18 stationary phase. Samples were fractionated using a linear gradient (mobile phase A: 0.1% trifluoroacetic acid; mobile phase B: acetonitrile) and elution was monitored by absorbance measured at three wavelengths (214 nm, 260 nm, 280 nm). Analyte peaks were grouped into 17 fractions defined by elution time (**Figure 2**). The fractions were lyophilized, resuspended in bath solution [the extracellular bath solution contained (mM): 140 NaCl, 3 KCl, 1 MgCl_2_, 1 CaCl_2_, 10 HEPES; pH was adjusted to 7.3 with NaOH] and tested for inhibitory activity against a recombinant Nav1.8 clone from grasshopper mice (*Onychomys torridus*, OtNav1.8).

**Figure 2.**
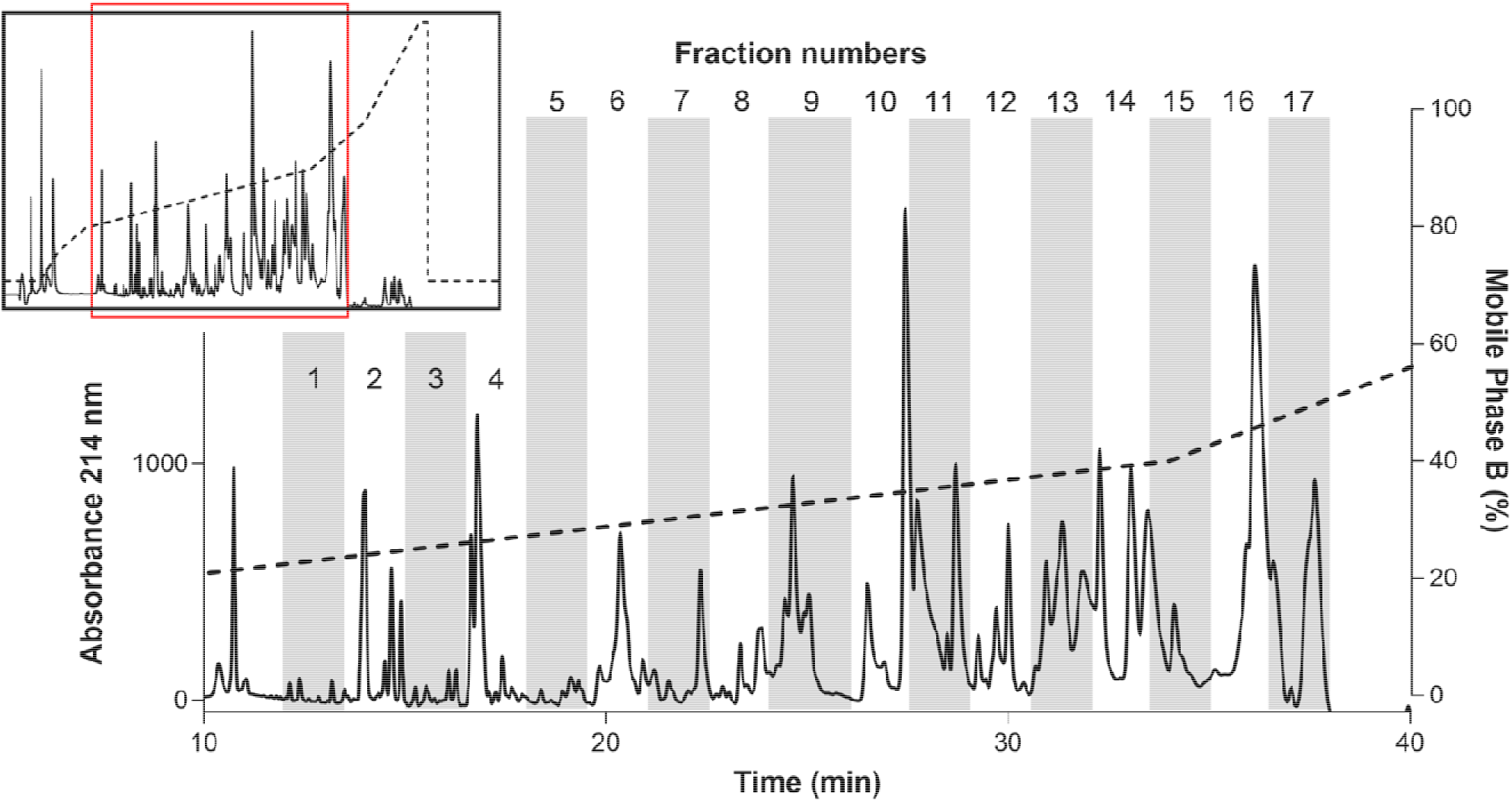
Chromatogram from Reversed Phase Liquid Chromatography (RPLC) fractionation of AZ bark scorpion venom. The inset shows the complete chromatogram. The main figure shows a magnification of the region indicated by the red box in the inset. The dashed line represents the % mobile phase B gradient. The peaks were divided into 17 fractions.

### 2.2. Identification of inhibitory fractions from AZ bark scorpion venom

Whole-cell patch clamp electrophysiology was used to measure the effects of venom and venom fractions on tetrodotoxin (TTX) resistant Na^+^ current recorded from OtNav1.8 expressed in the rodent neuroblastoma cell line (ND7/23 cells). Currents were elicited by a 100-msec depolarization to +20 mV from a holding potential of −80 mV before and after application of venom (1.2 μg/mL) and venom fractions. Pre- and post-current traces were compared to detect inhibition of Na^+^ current. While the venom inhibits OtNav1.8 activity completely, low concentrations (0.1 – 0.3 μg/mL) of fractions 7, 11, 12, 13, and 16 partially inhibited Nav1.8 TTX resistant Na^+^ current (**Figure 3A-L**). As shown in the summary graph in **Figure 3**, venom, F7, F11, F12, F13 and F16 significantly (* P < 0.05) inhibited OtNav1.8 Na^+^ current measured in pA/pF (venom: from 168.6 ± 10.187 to 5.084 ± 0.575, n = 6; F7: from 147.448 ± 18.149 to 80.770 ± 20.430, n = 5; F11: from 251.699 ± 18.683 to 98.224 ± 20.718, n = 6; F12: from 216.320 ± 34.260 to 120.493 ± 19.443, n = 6; F13: from 92.243 ± 20.231 to 37.364 ± 7.774, n = 5; and F16: from 129.713 ± 20.386 to 84.763 ± 18.208, n = 6). Fractions 1 – 6, 8 – 10, 14, 15 and 17 had no effect on OtNav1.8 Na^+^ current (Figure S1). Notably, with the exception of fraction 7, which eluted between 22 and 23 minutes, the majority of inhibitory fractions eluted between 30 and 37 minutes (**Figure 2**).

**Figure 3.**
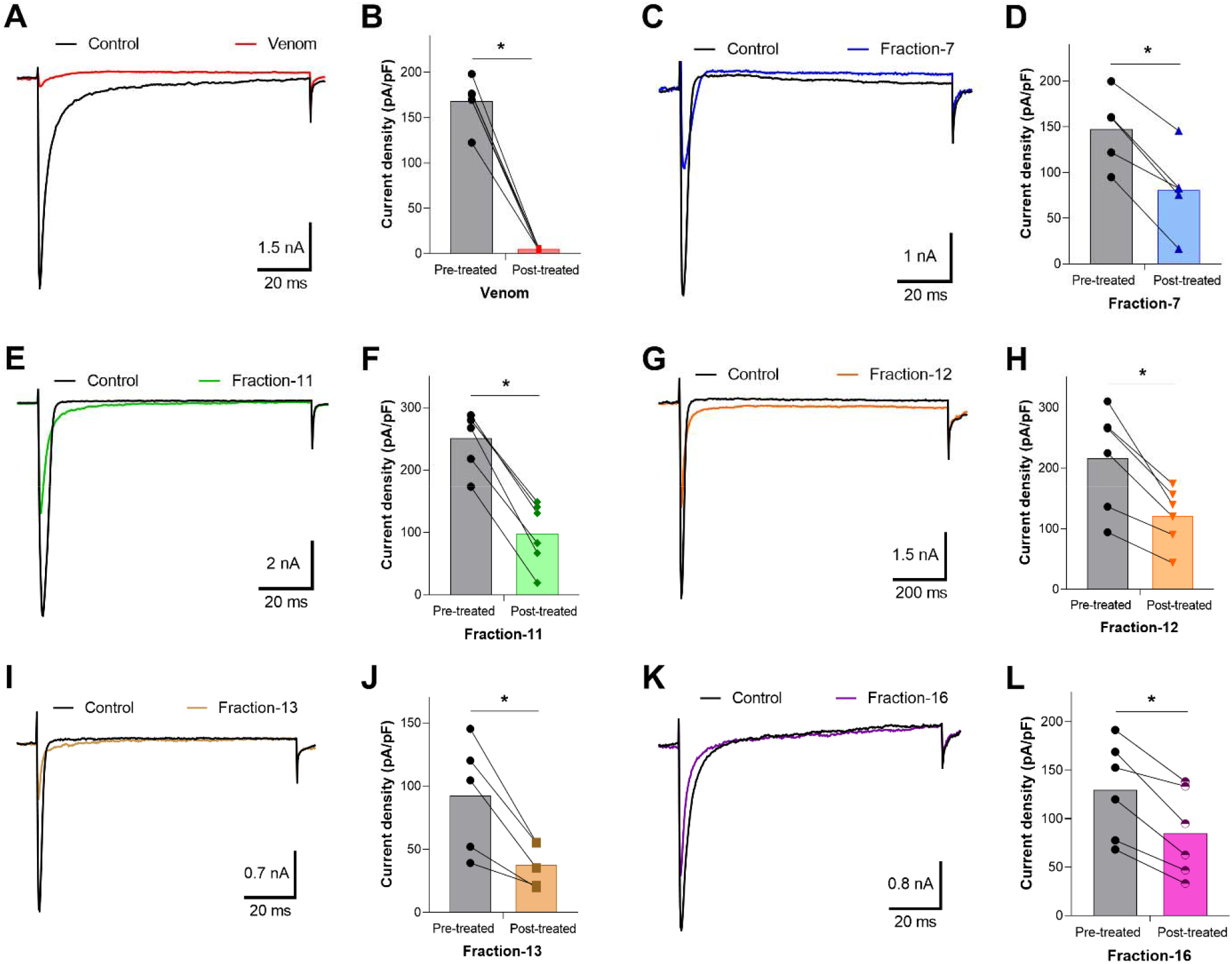
Effects of the venom and RPLC fractions on grasshopper mice recombinant OtNav1.8 Na^+^ current. (**A**) Representative OtNav1.8 currents before (black trace) and after (red trace) application of venom (1.2 μg/mL). The currents were elicited by a 100-msec depolarization to +20 mV from a holding potential of −80 mV before and after application of venom. (**B**) Summary graph of Na^+^ currents quantified in whole-cell voltage clamped ND7/23 cells transfected with OtNav1.8 before (black circles, gray bar) and after (red squares, red bar) application of venom. Summary data from experiments (n = 6 cells) identical to those shown in **A**. * P < 0.05 vs. before application of venom. (**C, E, G, I**, and **K**) Representative OtNav1.8 currents before (black traces) and after application of fractions-7 (blue trace), 11 (green trace), 12 (orange trace), 13 (light brown trace), and 16 (violet trace). The currents were elicited by a 100-msec depolarization to +20 mV from a holding potential of −80 mV before and after application of fractions (0.1 – 0.3 μg/mL) 7, 11, 12, 13, and 16. (**D, F, H, J**, and **L**) Summary graphs of Na^+^ currents quantified in whole-cell voltage clamped ND7/23 cells transfected with OtNav1.8 before (black circles, gray bars) and after application of fractions 7 (blue triangles, light blue bar), 11 (green diamonds, light green bar), 12 (orange triangles, light orange bar), 13 (brown circles, light brown bar), and 16 (violet and white circles, pink bar). Summary data from experiments (n = 5 – 6 cells) identical to those shown in **C, E, G, I**, and **K**. * P < 0.05 vs. before application of fractions-7, 11, 12, 13, and 16.

### 2.3. Ion exchange chromatography of inhibitory fractions-7, 11, 12 and 13 and identification of inhibitory subfractions

Samples of inhibitory fractions 7, 11, 12, and 13 were further fractionated into subfractions using ion exchange chromatography (IEC) and a linear increase in ionic strength using KCl (only data from fractions 7 and 11 are shown). Fraction 7 separated into 14 subfractions designated A – N (**Figure 4**). After desalting, samples of each subfraction were screened for inhibition of OtNav1.8 TTX-R Na^+^ current. Whole-cell patch clamp electrophysiology was used to measure the effects of venom subfractions on TTX-R Na^+^ current recorded from OtNav1.8 expressed in ND7/23 cells. Samples of subfractions were diluted in bath solution (0.1 – 0.3 μg/mL). Currents were elicited by a 100-msec depolarization to +20 mV from a holding potential of −80 mV before and after application of the venom or subfractions. Pre- and post-current traces were compared to detect inhibition of current. Subfractions 7A, 7C, 7E, 7F, 7M, and 7N inhibited OtNav1.8 Na^+^ current (**Figure 5A-L**). As shown in the summary graph in **Figure 5**, subfractions 7A, 7C, 7E, 7F, 7M, and 7N significantly (* P < 0.05) inhibited OtNav1.8 Na^+^ current measured in pA/pF (7A: from 105.559 ± 14.358 to 24.918 ± 4.720, n = 5; 7C: from 112.855 ± 11.219 to 59.633 ± 7.176, n = 7; 7E: from 236.654 ± 39.063 to 105.479 ± 22.385, n = 6; 7F: from 302.570 ± 22.213 to 22.956 ± 2.780, n = 7; 7M: from 208.440 ± 46.256 to 33.073 ± 9.577, n = 4; and 7N: from 426.745 ± 55.432 to 8.404 ± 1.536, n = 6).

**Figure 4.**
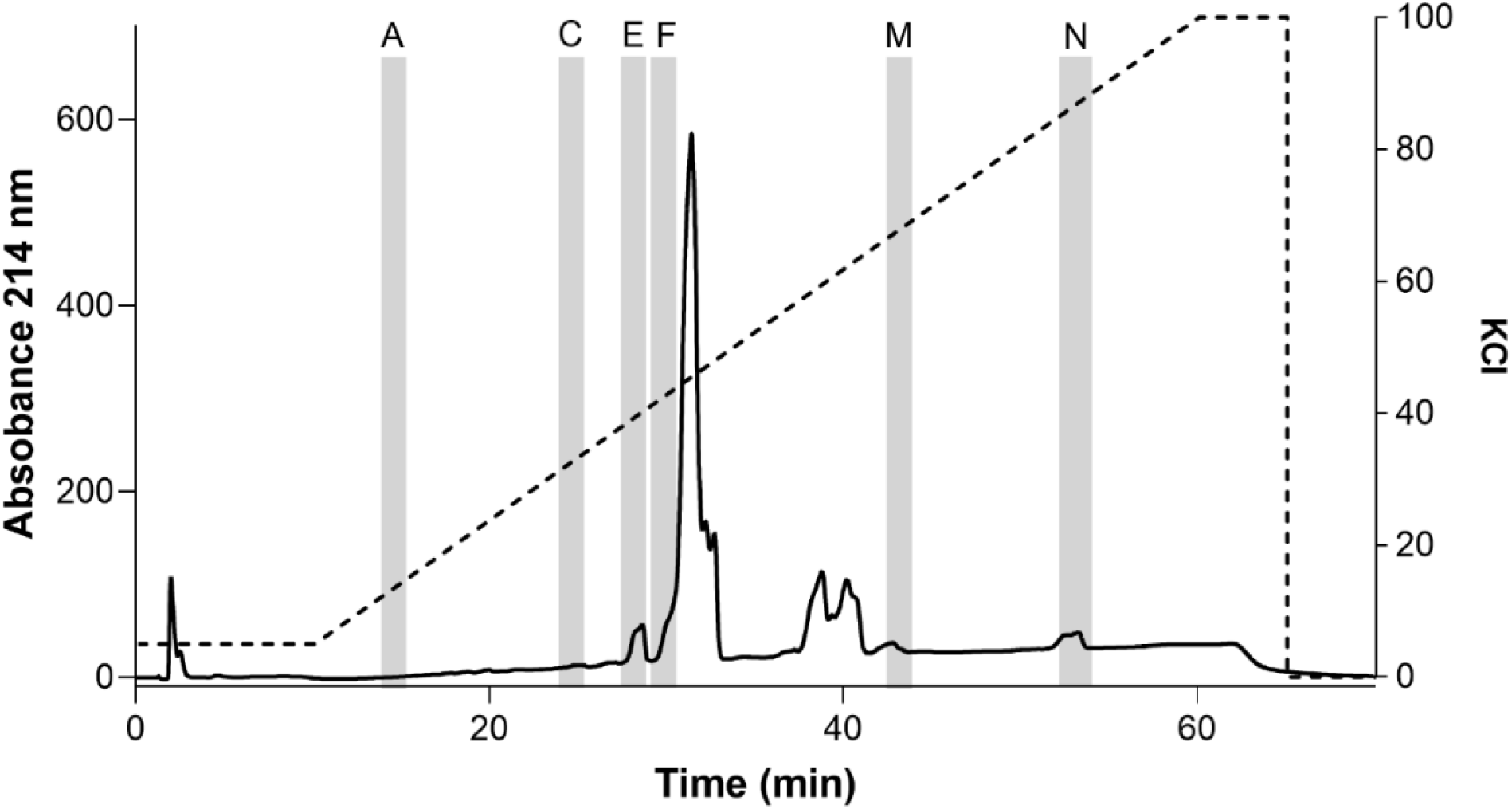
Ion Exchange Chromatography (IEC) sub-fractionation profile of fraction 7. Venom fraction 7 was further separated using an IEC column and a linear gradient of KCL buffer. Fraction 7 separated into 14 subfractions designated as A-N. Subfractions were tested against the recombinant OtNav1.8 clone for inhibitory activity. Subfractions A, C, E, F, M, and N (highlighted in the chromatogram with gray columns) inhibited TTX-R Na^+^ current recorded from OtNav1.8 expressed in ND7/23 cells. The dashed line shows the % gradient for mobile phase B (10mM KH_2_PO_4_, 0.5M KCl, 20% Acetonitrile, pH 2.7−3.0). Absorbance shown at wavelength 214 nm.

**Figure 5.**
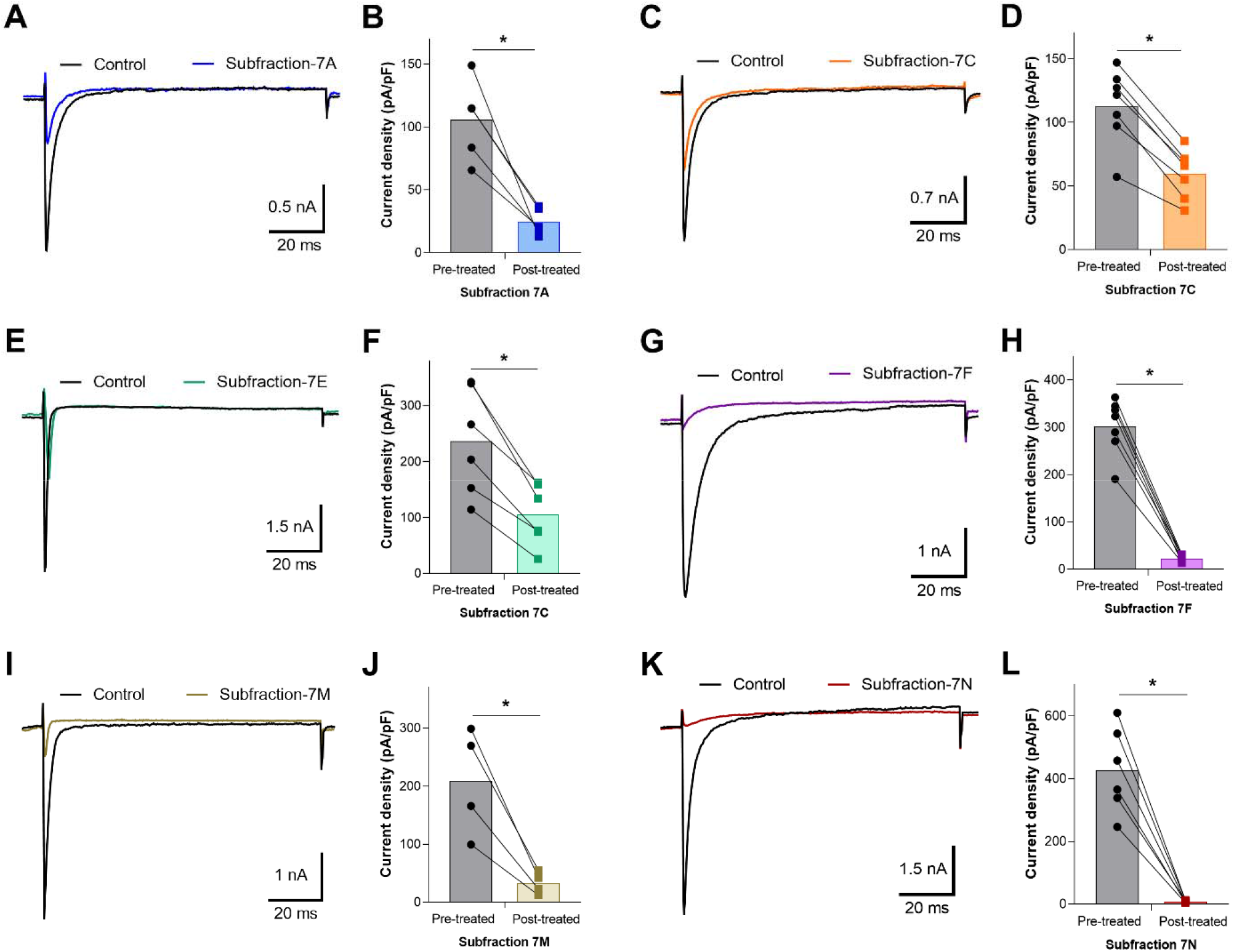
Effects of F7 subfractions on recombinant OtNav1.8 Na^+^ current. (**A, C, E, G, I**, and **K**) Representative OtNav1.8 currents before (black traces) and after application of 0.1 – 0.3 μg/mL of subfractions 7A (blue trace), 7C (orange trace), 7E (green trace), 7F (violet trace), 7M (cumin color trace), and 7N (brown trace). The currents were elicited by a 100-msec depolarization to +20 mV from a holding potential of −80 mV before and after application of subfractions 7A, 7C, 7E, 7F, 7M, and 7N. (**B, D, F, H, J**, and **L**) Summary graphs of Na^+^ currents quantified in whole-cell voltage clamped ND7/23 cells transfected with OtNav1.8 before (black circles, gray bars) and after application of subfractions 7A (blue squares, light blue bar), 7C (orange squares, light orange bar), 7E (green squares, light green bar), 7F (violet squares, light violet bar), 7M (cumin color squares, light cumin color bar), and 7N (red squares, red bar). Summary data from experiments (n = 4 – 7 cells) identical to those shown in **A, C, E, G, I**, and **K**. * P < 0.05 vs. before application of subfractions 7A, 7C, 7E, 7F, 7M, and 7N.

Fraction 11 separated into 10 subfractions designated as A – J (**Figure 6A**), only subfractions 11E, 11I and 11J inhibited OtNav1.8 Na^+^ current (**Figure 6B-G**). As shown in the summary graph in **Figure 6**, subfractions 11E, 11I, and 11J significantly (* P < 0.05) inhibited OtNav1.8 Na^+^ current measured in pA/pF (11E: from 82.530 ± 18.799 to 6.911 ± 0.919, n = 5; 11I: from 205.128 ± 30.427 to 89.581 ± 13.328, n = 7; and 11J: from 181.488 ± 25.314 to 63.843 ± 12.313, n = 6). Notably, none of the principal peaks from either fraction 7 or 11 inhibited OtNav1.8 Na^+^ current (**Figures S2 & S3**). Instead, the inhibitory subfractions consisted of minor peaks in both the fraction 7 and 11 profiles (highlighted in the chromatograms with gray columns) (**Figures 4 & 6A**). This suggests that inhibitory peptides are expressed in low quantity compared to other peptides and may not represent venom components that are key for prey capture and predator defense. A comparison of venom fractions with subfractions also suggests that the inhibitory effects are concentration dependent. For example, low concentrations (0.1 – 0.3 μg/mL) of fractions 7 and 11 did not inhibit OtNav1.8 activity completely (**Figure 3C & E**). However, similar concentrations (0.1 – 0.3 μg/mL) of subfractions 7F, 7M, 7N and 11E completely blocked OtNav1.8 activity (**Figures 5 & 6**).

**Figure 6.**
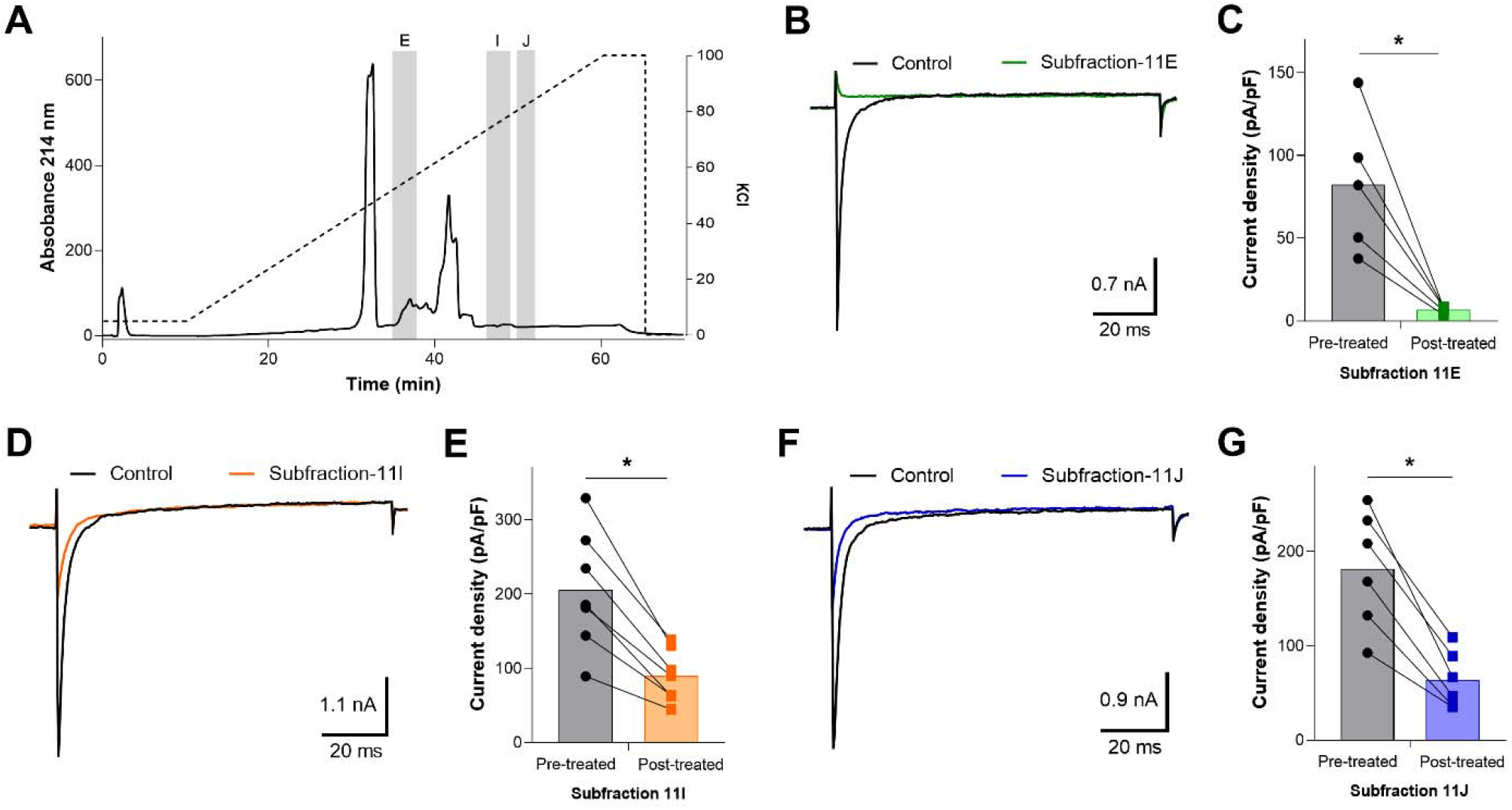
Effects of F11 subfractions on recombinant OtNav1.8 Na^+^ current. (**A**) Ion Exchange Chromatography (IEC) sub-fractionation profile of fraction 11. Venom fraction 11 was further separated using an IEC column and a linear gradient of KCL buffer. Fraction 11 separated into 10 subfractions designated as A – J. Subfractions were tested against the recombinant OtNav1.8 clone for inhibitory activity. Subfractions E, I, and J (highlighted in the chromatogram with gray columns) inhibited TTX-R Na^+^ current recorded from OtNav1.8 expressed in ND7/23 cells. The dashed line shows the KCl gradient used for elution of the compounds. Absorbance shown at wavelength 214 nm. (**B, D**, and **F**) Representative OtNav1.8 currents before (black traces) and after application of subfractions 11E (green trace), 11I (orange trace), and 11J (blue trace). The currents were elicited by a 100-msec depolarization to +20 mV from a holding potential of −80 mV before and after application of 0.1 – 0.3 μg/mL of subfractions 11E, 11I, and 11J. (**C, E**, and **G**) Summary graphs of Na+ currents quantified in whole-cell voltage clamped ND7/23 cells transfected with OtNav1.8 before (black circles, gray bars) and after application of subfractions 11E (green squares, light green bar), 11I (orange squares, light orange bar), and 11J (blue squares, light blue bar). Summary data from experiments (n = 5 – 7 cells) identical to those shown in **B, D**, and **F**. * P < 0.05 vs. before application of subfractions 11E, 11I, and 11J.

### 2.4. Electrophysiological characterization of voltage-dependent effects of AZ bark scorpion venom and inhibitory subfractions

We previously showed that AZ bark scorpion venom has no effect on grasshopper mice OtNav1.8 channel fast inactivation *[49]*. Therefore, we hypothesized that proteins may inhibit OtNav1.8 by binding to the extracellular side of the pore and blocking Na^+^ ion flux. Sodium ion channels are gated (opened, inactivated, closed) by changes in voltage across the cell membrane. Membrane voltage has no effect on inhibition induced by proteins or small molecules that occlude the pore. We reasoned that if proteins inhibit OtNav1.8 by blocking the pore, the inhibitory effect would be independent of voltage. Alternatively, proteins could bind extracellular channel loops and inhibit Nav1.8 through voltage dependent mechanisms. To address this, we tested the effects of venom (1.2 μg/mL) on OtNav1.8 held at different membrane potentials and found that the inhibitory effect of venom is voltage dependent. Hyperpolarized membrane potentials (e.g. –120 mV) reduced the inhibitory effects of the AZ bark scorpion venom on the channels (**Figure 7**). A comparison of the Na^+^ current (**Figure 7A, B** and **C**) with the corresponding current voltage (I-V) relationship (**Figure 7D**) shows a decrease in Na^+^ currents after application of the AZ bark scorpion venom followed by an increase in current after hyperpolarizing the cell membrane to –120 mV (i.e., reduction in the inhibitory effect of the venom). The total Na^+^ current at 10 mV decreased from −564.52 ± 74.21 pA/pF (control, n = 5) to −78.86 ± 61.57 (application of venom, n = 5). Hyperpolarization of the cell membrane to −120 mV for 30 sec induced a negative shift in the voltage dependence of activation from 10 mV (control) to −20 mV (hyperpolarization) (**Figure 7D**). The peak Na^+^ amplitude at −20 mV increased from −56.73 ± 29.18 pA/pF (application of venom, n = 5) to −295.89 ± 39.31 pA/pF (hyperpolarization).

**Figure 7.**
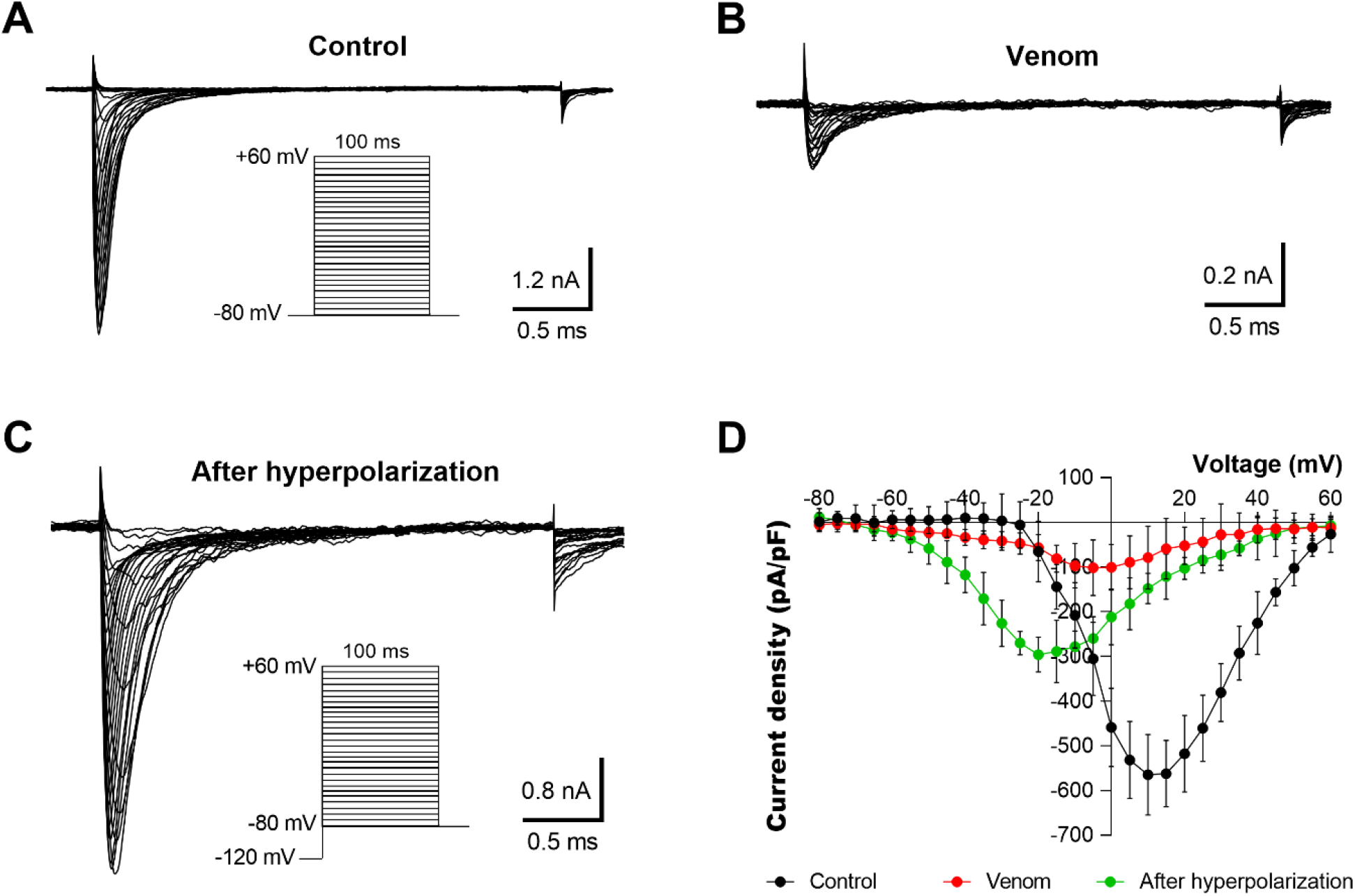
Inhibitory effects of AZ bark scorpion venom on OtNav1.8 channels are voltage dependent. (**A** and **B**) Representative activation current traces of OtNav1.8 channel expressed in ND7/23 cells in absence (control, **A**) and in presence (**B**) of venom (1.2 μg/mL). The protocol is shown in inset. Total currents were elicited by a family of 100 ms depolarization from −80 mV to +60 mV in 5 mV increments. (**C**) Representative activation current traces of OtNav1.8 in the presence of the venom after hyperpolarizing the cell membranes (the protocol is shown in inset). The current-voltage curve was generated by voltage-clamp protocols consisting of holding potential of −120 mV for 30 sec followed by a family of 100 ms depolarization from −80 mV to +60 mV in 5 mV increments. (**D**) I-V curves of the total Na^+^ currents in the absence (control, black circles) and presence of the venom (red circles) and after hyperpolarization (green circles) at a holding potential of −120 mV for 30 sec followed by a family of 100 ms depolarization from −80 mV to +60 mV in 5 mV increments. The inhibitory effects of venom on Na^+^ current were reduced when ND7/23 cells (expressing OtNav1.8) hyperpolarized to a holding potential of −120 mV compared to −80 mV. Summary data from experiments (n = 3 – 5 cells) identical to those shown in **A, B**, and **C**.

We also tested the F7 subfractions (7A, 7E, 7F, 7M, and 7N) that inhibited OtNav1.8 current and found that the inhibitory effects of the subfractions could be reduced by hyperpolarizing the cell membranes (**Figure 8**, data shown for subfraction 7A only while subfractions 7F, 7M, and 7N shown in **Figures S4, S5 & S6**). A comparison of the Na^+^ current (**Figure 8A, B** and **C**) with the corresponding current voltage (I-V) relationship (**Figure 8D**) shows a decrease in Na^+^ currents after application of the subfraction 7A followed by an increase in current after hyperpolarizing the cell membrane to –120 mV. The total Na^+^ current at 10 mV decreased from −982.05 ± 130.12 pA/pF (control, n = 4) to −315.52 ± 70.08 (application of subfraction 7A, n = 4). Hyperpolarization of the cell membrane to −120 mV for 30 sec induced a negative shift in the voltage dependence of inactivation from 10 mV (control) to −15 mV (hyperpolarization) (**Figure 8D**). The peak Na^+^ amplitude at −15 mV increased from −239.65 ± 31.81 pA/pF (application of subfraction 7A, n = 4) to −786.28 ± 92.07 pA/pF (hyperpolarization). These results suggest that bark scorpion venom proteins may inhibit grasshopper mice Nav1.8 Na^+^ currents by modulating voltage dependent gating mechanisms.

**Figure 8.**
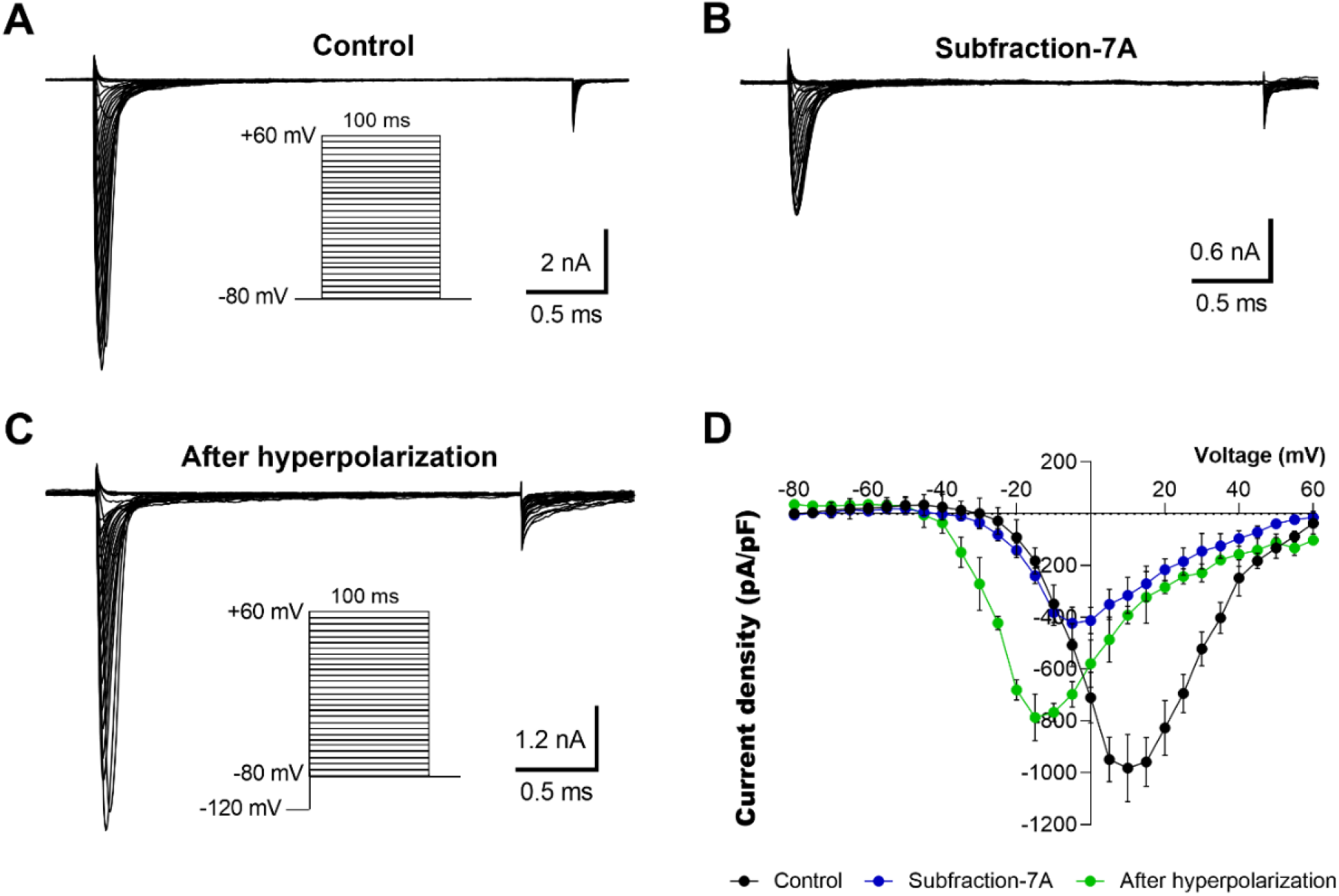
Inhibitory effects of subfraction 7A on OtNav1.8 channels are voltage dependent. (**A** and **B**) Representative activation current traces of OtNav1.8 channel expressed in ND7/23 cells in absence (control, **A**) in presence (**B**) of the subfraction 7A (0.1 – 0.3 μg/mL). The protocol is shown in inset. Total currents were elicited by a family of 100 ms depolarization from −80 mV to +60 mV in 5 mV increments. (**C**) Representative activation current traces of OtNav1.8 in presence of the subfraction 7A after hyperpolarizing the cell membranes (the protocol is shown in inset). The current-voltage curve was generated by voltage-clamp protocols consisting of holding potential of −120 mV for 30 sec followed a family of 100 ms depolarization from −80 mV to +60 mV in 5 mV increments. (**D**) I-V curves of the total Na^+^ currents in the absence (control, black circles) and presence of the subfraction 7A (blue circles) and after hyperpolarization (green circles) at a holding potential of −120 mV for 30 sec followed by a family of 100 ms depolarization from −80 mV to +60 mV in 5 mV increments. The inhibitory effects of subfraction 7A on Na^+^ current were reduced when ND7/23 cells (expressing OtNav1.8) were hyperpolarized to a holding potential of −120 mV compared to −80 mV. Summary data from experiments (n = 3 – 5 cells) identical to those shown in **A, B**, and **C**.

### 2.5. Proteomic analyses of AZ bark scorpion venom subfractions that inhibit OtNav1.8 Na^+^ current

Mass spectrometry-based bottom-up proteomic analyses were used to determine the primary structure of proteins from inhibitory subfractions of F7 and F11. Due to insufficient protein in a number of subfractions, only F7 C and F11 E and I provided proteomic results. In total, nine proteins were identified using multiple database search engines from the subfractions that showed bioactivity against OtNav1.8; four of these proteins were unique to the bioactive subfractions F11E (**Table 1**). The proteomic analysis also led to the identification of 18 proteins belonging to subfractions that did not show bioactivity, of which 13 were exclusively found in non-bioactive subfractions (summary of results provided in **Table S1**). Interestingly, each analyzed subfraction contained multiple proteins active against sodium and potassium channels along with other venom proteins such as enzymes. Typically, the proteins within a single subfraction demonstrated some degree of similarity in their sequences and could be grouped accordingly. As an example, **Table 1** provides the sequence comparison between four of the six identified proteins from subfraction 11E. Sequences of the four proteins, NaTx-22, NaTx-4, NATx-36, and NaTx-13, were identified by searching bottom-up proteomic data against a transcriptome database of the venom glands of Arizona bark scorpion (provided by D.R. Rokyta, Florida State University). The portions of sequences of NaTx-22, NaTx-4, NaTx-36, and NaTx-13 that correspond to mass spectrometry-identified tryptic peptides are underlined in black (**Table 1**). All four proteins exhibited the presence of multiple cysteine and lysine residues characteristic of scorpion venom toxins that bind voltage-gated sodium channels. The four proteins also have a “CXCE” motif where X = an aromatic residue (tryptophan, tyrosine or phenylalanine). Three of the toxins have multiple prolines in the C-terminal end. Additionally, two of the toxins exhibit a “PTXPIP” motif where X = tyrosine or phenylalanine. The other proteins identified in subfraction 11E are bNaTx-7 and VP-16. While bNaTx-7 shares a similar abundance of lysines and prolines along with the “CXCE” motif of the four toxins in **Table 1**, it was also identified in a non-bioactive subfraction and thus removed from consideration. VP-16 is a larger protein than the others (10.7 kDa compared to approximately 7 kDa for the other four). It has a similar abundance of prolines and lysines but lacks either the “CXCE” or “PTXPIP” found in the others. Like bNaTx-7, VP-16 was also identified in a non-bioactive subfraction and removed from consideration.

**Table 1.**
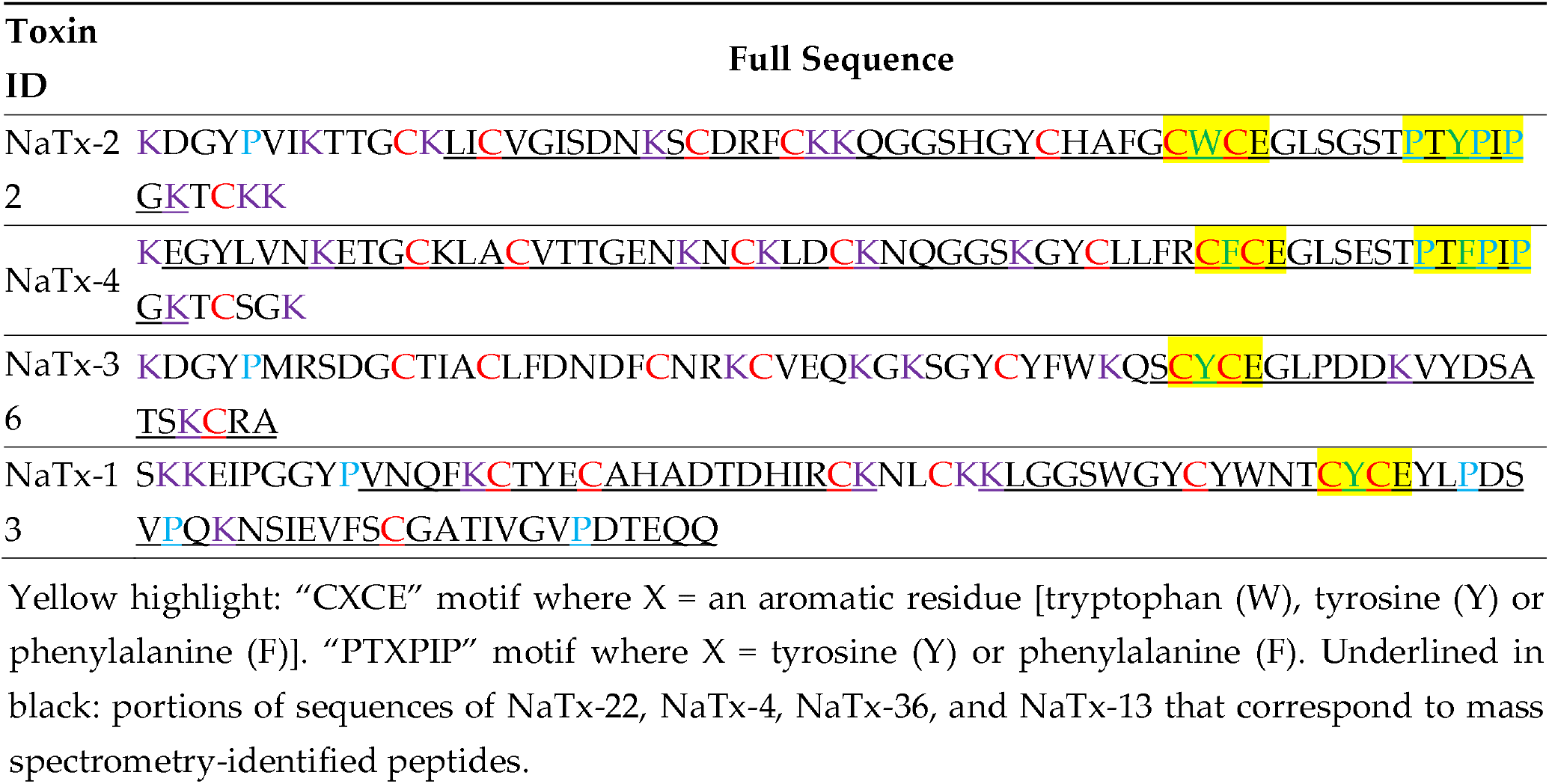
Primary structure of toxin proteins identified from subfraction 11E that inhibited OtNav1.8 activity.

### 2.6. Structural characterization of NaTx-22 from subfraction 11E that inhibits OtNav1.8

Proteomic analyses of AZ bark scorpion venom subfractions that inhibited grasshopper mice Nav1.8 Na^+^ current identified four novel toxin proteins with a mass and primary sequence between 40 and 65% conserved with toxins previously isolated from scorpion venoms. Importantly, the toxin proteins were unambiguously characterized through the identification of unique tryptic peptides as shown in **Figure 9A** for NaTx-22 and in **Figures S7A, S8A**, and **S9A** for the other three proteins presented in **Table 1**. In order to determine the possible presence of similar proteins in other related scorpion species, the sequences from the four proteins were searched against other proteins using the National Center for Biotechnology Information Protein Basic Local Alignment Search Tool, which produced a match with other scorpion toxins (NCBI Protein BLAST, **Figures 9B, S7B, S8B**, and **S9B**). NaTx-22 has sequence identity of 65% compared to Cex5, a toxin cloned from *Centruroides exilicauda* venom gland (*Centruroides exilicauda* is now renamed as *Centruroides sculpturatus*; UniProtKB/Swiss-Prot: Q68PH0.1). In addition, NaTx-22 is between 62 and 63% conserved with two toxins identified from *C. noxius* venom. NaTx-22 is predicted to be a *β*-toxin based on primary structure. The proposed disulfide bridge arrangement of NaTx-22 is C1-C8, C2-C5, C3-C6, and C4-C7 (based on homology with Cex5, UniProtKB/Swiss-Prot: Q68PH0.1, Figure 9C) [60]. NaTx-22 has a theoretical molecular weight of 6962.08 Da (ExPASy ProtParam, http://web.expasy.org/protparam). The SWISS-MODEL (http://swissmodel.expasy.org/) was used to propose the 3D-structure of NaTx-22 and the other three toxins presented in **Table 1** (**Figures 9D, S7D, S8D**, and **S9D**).

**Figure 9.**
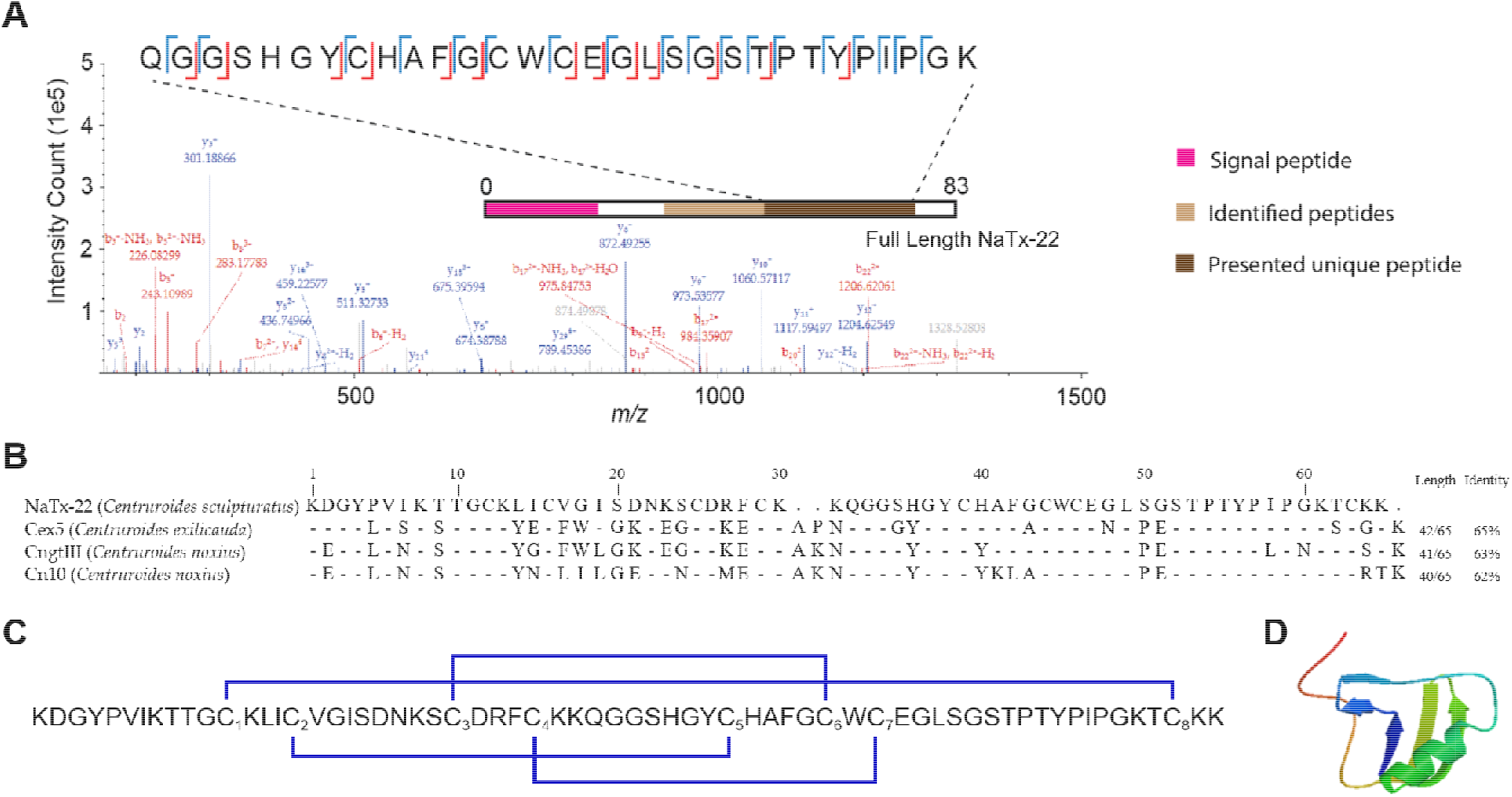
Primary structure of NaTx-22, identified from AZ bark scorpion venom that inhibit Nav1.8 Na^+^ current. (**A**) Tandem mass spectrometry-based identification of a unique peptide from NaTx-22. Experimental fragment ions matched against in silico-generated fragment ions (using a 0.5 Da mass tolerance) are indicated in red (b-ions, containing the N-terminus) or blue (y-ions, containing the C-terminus). Identified cleavage sites are recapitulated in the inset showing the exact sequence of the peptide. The position of the identified tryptic peptide within the whole sequence of NaTx-22 is indicated in brown on the bar representing the full sequence of the toxin protein. This tryptic peptide has a sequence unique to this toxin, and the intact mass of the peptide was matched with a 5.3 ppm deviation over the theoretical value. (**B**) Sequence alignment of NaTx-22 with homology toxins from NCBI BLAST. UniProt Knowledgebase (UniProtKB)/Swiss-Prot accession codes for these retrieved sequences are Q68PH0.1 (Cex5), P45664.1 (CngtIII), and Q94435.2 (Cn10). Hyphen-minus represents identical amino acid residues, and dots indicate the lack of residue at the position. The toxin lengths and percentages of sequence identities are given on the right. (**C**) Disulfide bridge arrangement (in blue) of NaTx-22 proposed by homology with Cex5. Cysteines were numbered by their order of appearance in the sequence. (**D**) SWISS-MODEL (http://swissmodel.expasy.org/) proposed the 3D-structure of NaTx-22.

## 3. Discussion

Chronic pain causes human suffering and a loss of economic productivity. Related opioid abuse imposes a burden on community health systems. A better understanding of the mechanisms underlying transmission of pain signals, particularly mechanisms that inactivate or block signals would advance efforts to develop drugs that block pain without addiction [61]. Pain pathway neurons transmit pain signals to the brain via action potentials. Nav1.8 generates the majority of Na^+^ current underlying the action potentials that carry pain signals to the brain. Tissue damage due to injury, aging or disease causes biochemical changes in neurons that activate Nav.18 to initiate action potentials [1,5–8]. Nav1.8 has two inactivation mechanisms that prevent the flux of Na^+^ ions into nociceptive neurons, blocking the action potentials that carry pain signals to the brain. Fast inactivation blocks Na^+^ flux for milliseconds, while slow inactivation blocks channels over longer timescales. The biophysical mechanisms that govern activation and inactivation gating provide a strategy for developing non-addictive pain treatments.

Animal venoms are a rich source of proteins that target pain-pathway neurons [24–30]. Venom proteins modulate the voltage-gated sodium channels that regulate transmission of pain signals to the brain, thus, providing a toolkit for examining the biophysical mechanisms that inhibit channel activity and block pain signal transmission [62]. Moreover, proteins that modulate VGSC activity can serve as templates for structure-guided engineering of drugs that block pain [37–40]. However, examining Nav1.8 inactivation mechanisms has been challenging. While Nav1.8 is preferentially expressed in nociceptive neurons where it plays a role in neuropathic and inflammatory pain, highlighting the potential for Nav1.8 to serve as an alternative drug target to Nav1.7, the biophysical mechanisms that regulate Nav1.8 inactivation are not completely understood [7,8,10–18]. Progress has been hindered by a lack of venom proteins that modulate Nav1.8 inactivation. Zhang *et al.* [59] screened venoms from multiple species and identified only one protein from snake venom that inhibits Nav1.8 currents expressed in rat DRG. The three-finger protein μ-EPTX-Na1a, isolated from the Chinese cobra (*Naja atra*), inhibits Nav1.8 through a depolarizing shift of activation and a repolarizing shift of inactivation [59].

AZ bark scorpions (*Centruroides sculpturatus*) produce venom proteins that reversibly bind VGSCs [50–56]. Grasshopper mice (*Onychomys torridus*) feed on AZ bark scorpions but show little response when stung by scorpions [49,57,58]. When mice are stung by scorpions, the venom inhibits Nav1.8 activity, blocking the compound action potentials that carry pain signals to the brain – the venom acts as an analgesic to block pain in these mice (**Figure 10**) [49]. Whole cell patch clamp recordings showed that venom inhibited Nav1.8 currents and blocked action potentials in nociceptive neurons from mice [49]. Injections of venom into the hind paw of mice blocked paw-licking behavior in response to injections of the pain inducing chemical formalin. Thus, grasshopper mice provide a system for characterizing peptides that inhibit Nav1.8 activity and block pain signal transmission. If we could determine how venom proteins bind mouse Nav1.8 and block pain signal transmission, we could apply that knowledge to develop proteins that would block pain in humans.

**Figure 10.**
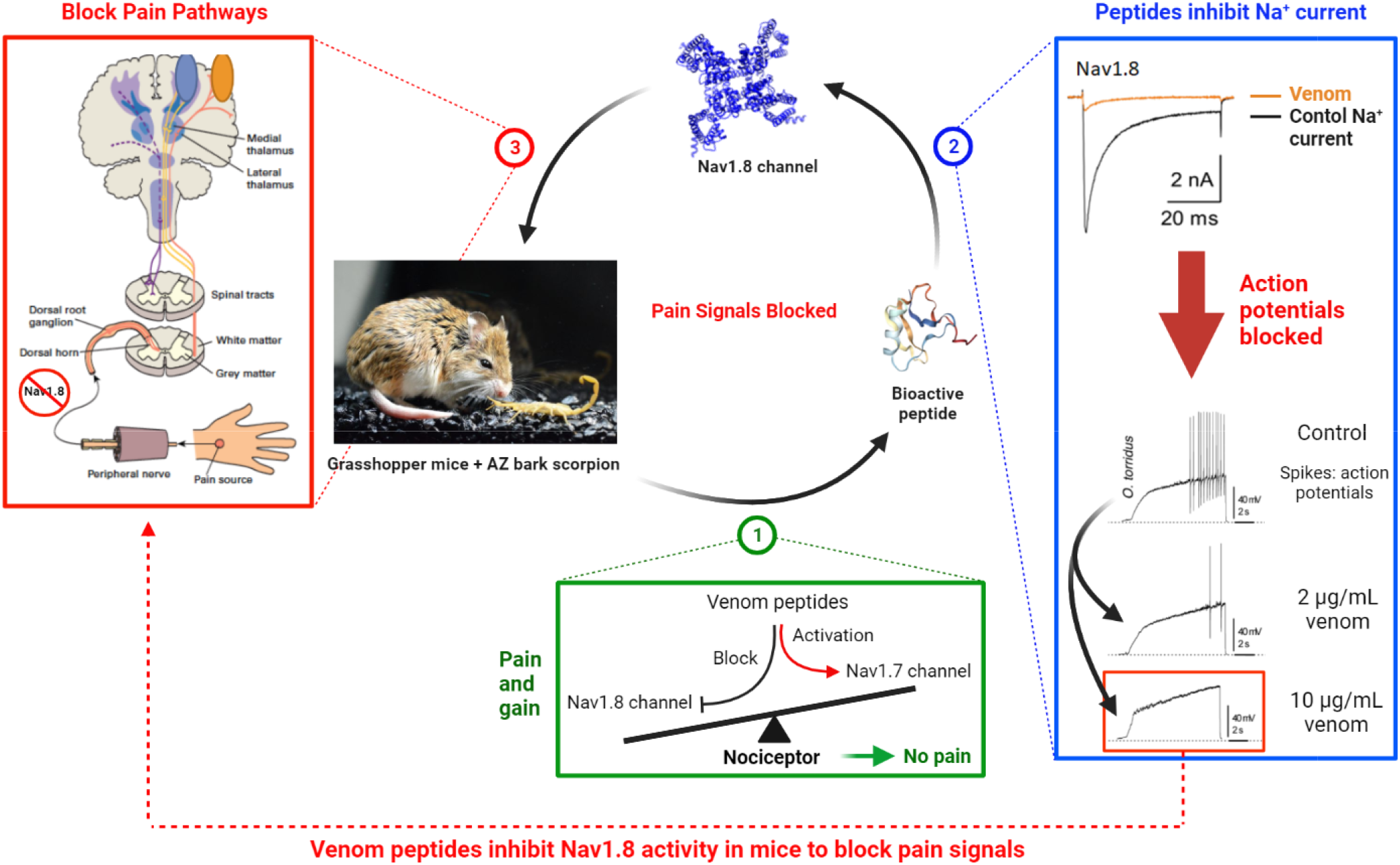
AZ bark scorpion venom blocks transmission of pain signals in grasshopper mice. When grasshopper mice are stung by AZ bark scorpions, the venom proteins inhibit Na^+^ ion flux through Nav1.8. Inhibition of Nav1.8 Na^+^ current blocks the generation of action potentials in the nociceptive neurons that carry pain signals to the brain. **Figure** modified from Rowe *et al*. [49]. Created with BioRender.com

Using a bidimensional (RPLC, IEC) fractionation/subfractionation and bioassay pipeline we isolated several venom subfractions from AZ bark scorpions that inhibit grasshopper mice Nav1.8 TTX resistant Na^+^ current. While the venom completely blocks TTX resistant Na^+^ current in our recombinant OtNav1.8 bioassay, many of the venom fractions only partially inhibited the Na^+^ current at low concentrations (0.1 – 0.3 μg/mL). However, similar concentrations (0.1 – 0.3 μg/mL) of more pure venom subfractions almost completely inhibited OtNav1.8. These results suggest the inhibitory effects of venom proteins are concentration dependent. It is also possible that individual toxin proteins vary in their inhibitory efficacy. Alternatively, less pure fractions may contain proteins that enhance Na^+^ current and oppose the effects of inhibitory proteins. Current work is focused on isolating and testing individual toxin proteins to determine the EC_50_ for each toxin.

There are a number of mechanisms that could inhibit VGSC Na^+^ current: 1) physical occlusion of the pore; 2) inhibition of activation by trapping the DII voltage sensor in the deactivated position; 3) fast inactivation via modulation of the DIV voltage sensor to engage the hinged lid on the intracellular side of the pore; and 3) slow inactivation via conformational change of the helices that line the pore. Spiders, cone snails, and scorpions produce venom proteins that reversibly bind Na^+^ channels [24,41,42]. While cone snail μ-conotoxins occlude the pore from the extracellular side [63], scorpion and spider proteins function as gating-modifiers by docking to the voltage sensors [29,30,45,64]. Scorpion α toxins immobilize the DIV voltage sensor in the inward position (resting state) to impair channel fast inactivation [64]. Immobilization of the DIV voltage sensor prevents the hinged lid from moving into the intracellular side of the pore. The *β* toxins trap the DII voltage sensor in the outward position to activate (open) the channel [45]. The snake venom toxin μ-EPTX-Na1a from the Chinese cobra inhibits Nav1.8 in rat DRG through a novel mechanism – the toxin causes a depolarizing shift in the voltage dependence of activation and a repolarizing shift in the voltage dependence of inactivation. Electrophysiological data from our previous work showed that AZ bark scorpion venom did not inhibit Nav1.8 Na^+^ current by engaging the fast inactivation hinged lid [49]. The electrophysiological characterization of venom subfractions from F7 in this study showed that the inhibitory effects of F7 subfractions could be reduced or removed by hyperpolarizing the cell membranes. Moreover, the hyperpolarizing holding potentials revealed that inhibitory subfractions from F7 negatively shift the voltage-dependence of OtNav1.8 activation. Taken together, these results suggest two potential mechanisms: 1) venom proteins may trap an OtNav1.8 voltage sensor(s) in the activated position to open channels at or near resting membrane potentials; and 2) proteins may function as gating modifiers to inhibit OtNav1.8 Na^+^ currents as opposed to pore blockers. Given that strong hyperpolarization of cell membranes can cause VGSCs to transition out of a slow inactivated state, the voltage dependent effects of the inhibitory F7 subfractions hint that venom proteins may inhibit Nav1.8 by inducing the channel to transition into a slow inactivated state. Future work should provide a detailed characterization of the electrophysiological effects of inhibitory toxin proteins from both F7 and F11 on the slow inactivation properties of grasshopper mice Nav1.8.

We identified the primary structure of four toxin proteins from a subfraction (F11E) of AZ bark scorpion venom that inhibits grasshopper mice Nav1.8. These four proteins have a primary structure, mass, cysteine-cysteine disulfide binding pattern and three-dimensional structure similar to Na^+^ channel toxins that have been previously described from scorpion venom (NCBI BLAST, ExPASy ProtParam, SWISS-MODEL). In particular, the four toxin proteins identified from AZ bark scorpion (*Centruroides sculpturatus*) venom were similar to toxins described from other *Centruroides* species (e.g., *C. noxius, C. suffusus suffusus, C. limpidus,* and *C. tecomanus*). In addition, one toxin protein from *C. sculpturatus* was similar to toxin proteins described from *Parabuthus transvalicus, Androctonus australis,* and *Tityus serrulatus* (**Figure S9**). Interestingly, several of the toxins identified as most similar to our candidate toxin proteins are either predicted or functionally confirmed to be *β* toxins. In general, *β* toxins function by trapping the domain II voltage sensor of VGSCs in the activated position, causing the channel to open at or near resting membrane potential. Results from this study suggest that inhibitory subfractions contain proteins that function as *β* toxins. Future work should determine the binding sites between grasshopper mice Nav1.8 and inhibitory toxin proteins, and whether the toxins trap a voltage sensor in the activated position.

Pain is necessary for survival because it warns of potential tissue damage, prompting individuals to seek medical attention. However, chronic pain due to injury, disease, aging or genetic disorders causes human suffering without providing pertinent medical information. Current drugs are not always adequate for managing pain and they may be addictive. Characterizing the structure-activity relationships between inhibitory proteins and OtNav1.8 will increase our understanding of the structural basis for Nav1.8 activation and inactivation gating. Moreover, amino acids at the core of structure-activity relationships are key to using a modified protein mutagenesis and screening approach similar to what has been accomplished engineering spider toxins that inhibit human Nav1.7 [39,65]. A better understanding of the molecular and biophysical mechanisms that regulate pain signal transmission, particularly mechanisms that inactivate signals, would advance efforts to develop drugs that block pain without addiction [61].

## 4. Conclusions

The voltage-gated sodium channel Nav1.8 is a potential target for neuropathic and inflammatory therapeutics. Yet, the biophysical mechanisms that regulate Nav1.8 gating are not completely understood. Arizona bark scorpion (*Centruroides sculpturatus*) venom proteins inhibit Nav1.8 and block pain in grasshopper mice (*Onychomys torridus*). Using liquid chromatography (reversed phase and ion exchange), a recombinant Nav1.8 bio-activity assay, and mass spectrometry-based bottom-up proteomics, we identified the mass and primary structure of four novel proteins from venom subfractions that inhibit Nav1.8 Na^+^ current. The proteins are between 40 and 60% conserved with Na^+^ channel toxins described from the venom of scorpions in the genera *Centruroides, Parabuthus, Androctonus, and Tityus*. Electrophysiology data showed that the inhibitory effects of bioactive subfractions can be removed by hyperpolarizing the channels, hinting that proteins may function as gating modifiers. These proteins provide a toolkit for investigating Nav1.8 gating mechanisms.

## 5. Materials and Methods

### 5.1. Venom extraction

AZ bark scorpions were collected from the Santa Rita Experimental Range, (University of Arizona, Santa Rita Mountains, AZ, USA). Crude venom was extracted from venom glands using electrical stimulation according to previously published protocols [27,49]. Crude venom samples were hydrated in sterile water, centrifuged, and filtered (0.45 μm sterile filter) to remove insoluble components. Aliquots of the supernatant (hereafter referred to as venom) were lyophilized and stored at −80° C.

### 5.2. Reversed Phase LC fractionation of the AZ bark scorpion venom

Lyophilized venom samples were hydrated in trifluoroacetic acid/distilled H_2_O (dH_2_O) 0.1/99.9 (*v/v*) to a final concentration of 0.0023 g/mL. Aliquots of 200 μL were injected into an InifinityLab Poroshell 120 Reverse Phase column (3.0 x 150 mm, 2.7 μm) and fractioned using an Agilent 1260 Infinity II High Performance Liquid Chromatography (HPLC) system (Agilent, Wilmington, DE, USA). Mobile phases were prepared as follows: mobile phase A, 0.1% trifluoroacetic (*v/v*) acid in double distilled, sterile water; mobile phase B, LC-MS grade acetonitrile. The separation was performed at a constant flow rate of 400 μL/min according to the following linear gradient of mobile phase B: 2% for 4 minutes; 12% at minute 6; 19.8% at minute 9; 39.2% by minute 34; 56% by minute 40; a ramp to 90% by minute 46; 1 minute of wash; decrease to 2% in 2 minutes (at minute 49); full re-equilibration until minute 53. The fractions were collected according to the following time scheme and were kept at 4° C; minutes 0-12, no fractions collected; 12-24, one fraction collected every 1.5 minutes; minutes 24-26, 1 fraction collected; 26-38, one fraction collected every 1.5 minutes; 38-53, no fractions collected. Venom elution was monitored using three wavelengths (214 nm, 260 nm, and 280 nm). Fractions were lyophilized and stored at −20° C.

### 5.3. Ion Exchange Liquid Chromatography to isolate individual peptides

Inhibitory fractions (F7, F11, F12, and F13) were further separated into subfractions using ion exchange liquid chromatography (IEC). Pooled samples of fractions were rehydrated to 600 μL with LC grade water and injected into a PolySULFOETHYL A column (200 x 2.1 mm, 5 μm particle size, 1000 Å pore size, PolyLC Inc., Columbia, MD, USA). A binary linear gradient [mobile phase A: 10 mM KH_2_PO_4_, 20% acetonitrile (v/v), pH 2.7−3.0; mobile phase B: 10 mM KH_2_PO_4_, 0.5 M KCl, 20% acetonitrile (v/v), pH 2.7−3.0] executed at a flow rate of 300 μL/min was used to subfraction samples using the following method: minute 0−5, 0% mobile phase B; minutes 5-50 linear ramp to 100% mobile phase B; minutes 50-55, 100% B ; minutes 55-56, mobile phase B was decreased to 0%. Re-equilibration in 100% mobile phase A was performed until minute 71. The subfraction collection was determined experimentally for each fraction based on the elution profile and locations of peaks in each fraction. Typically, fractions were collected for 2-3 minutes. The absorbance was measured at 214 nm, 230 nm, and 280 nm.

### 5.4. Culture and transfection of ND7/23 cells

Venom fractions and subfractions were screened for inhibitory activity against a recombinant Nav1.8 clone from grasshopper mice. The gene encoding *O. torridus* Nav1.8 was inserted into a plasmid with a fluorescent marker (pcDNA3.1-EGFP) for expression in a hybrid cell line (ND7/23). The recombinant Nav1.8 clone is referred to as OtNav1.8. ND7/23 cells were purchased from ATCC (American Type Culture Collection, ATCC, Manassas, VA, USA) and cultured under standard culture conditions (37° C in a humidified incubator supplying 5% CO_2_) in Dulbecco’s modified Eagle’s medium supplemented with 10% fetal bovine serum and 1% penicillin-streptomycin. For patch clamp recording (see below), ND7/23 cells were plated on cover glass chips treated with 0.01% Poly-L-lysine (Sigma, St. Louis, MO, USA). Plated cells were transfected with plasmids encoding α-, β1-, and β2-OtNav1.8 subunits genetically linked to NH2-terminal eGFP using Lipofectamine 3000 reagent (L3000015, Invitrogen, Carlsbad, CA, USA) as described in the manufacturer’s protocol. In brief, 60% confluent cells in a 35-mm dish were treated with 14 μg of total plasmid cDNA for 24-48 hr. Cells exhibiting green fluorescence were used for patch-clamp recording.

### 5.5. Electrophysiology recording

Whole-cell patch clamp electrophysiology is used to record the effects of venom fractions on OtNav1.8 Na^+^ current, expressed in ND7/23 cells. The venom and venom fractions are diluted in bath solution [the bath (extracellular) solution contained (mM): 140 NaCl, 3 KCl, 1 MgCl_2_, 1 CaCl_2_, 10 HEPES; pH was adjusted to 7.3 with NaOH] to the desired final concentration. The whole cell membrane currents were recorded using a low noise patch-clamp amplifier (Axopatch 200B; Molecular Devices, San Jose, CA, USA) interfaced via a Digidata 1550B (Molecular Devices) to a PC running the pClamp 11 suite of software (Molecular Devices). All currents were filtered at 1 kHz. Patch pipettes were pulled from borosilicate glass capillaries (World Precision Instruments, Inc., Florida, USA) using a model P-97 Flaming-Brown micropipette puller (Sutter Instrument) and fire-polished on a micro-forge (MF-830; Narishige Scientific Instrument, Japan) to have a 3–5 MΩ when backfilled with the pipette (intracellular) solution. The pipette solution contained (mM): 140 CsF, 10 NaCl, 1.1 EGTA, 10 HEPES; pH was adjusted to 7.3 with CsOH. Current traces were evoked by a 100 ms depolarizing potential of +20 mV from holding potential at −80 mV. While current-voltage curve was generated by voltage-clamp protocols consisting of holding potential of −80 mV followed by a family of 100 ms depolarization from −80 mV to +60 mV in 5 mV increments. In another set of experiments, current-voltage curve was generated by voltage-clamp protocols consisting of holding potential of −120 mV for 30 sec followed by a family of 100 ms depolarization from −80 mV to +60 mV in 5 mV increments. Tetrodotoxin (TTX; 500 nM) was added to the bath solution when recording recombinant OtNav1.8 currents from ND7/23 cells. All peptide effects were compared to baseline values obtained in vehicle (bath solution) in the same cell. After control responses were obtained, 1-10 μg/mL of scorpion venom or fraction peptides was added to the chamber (250 μl total volume) and mixed thoroughly, after which the current-clamp protocols were repeated. Experimental data were collected and analyzed with Clampfit 11 software (Molecular Devices).

### 5.6. Sample preparation for proteomic analyses

Subfractions were desalted using C18 spin columns (Thermo Fisher Scientific, Waltham, MA, USA), quantified by NanoDrop One (Thermo Fisher Scientific), and vacuum dried. Subfractions were then resuspended in 8 M urea buffered to pH 7.5 with 50 mM Tris-HCl before reduction and alkylation steps (using dithiothreitol and iodoacetamide, respectively). Trypsin digestion (Sequencing Grade Modified Trypsin, Promega, Madison, WI, USA) followed at 37° C for 18 h. Proteolytic peptides were purified using C18 ZipTips (Millipore Sigma, Burlington, MA, USA) and vacuum dried.

### 5.7. Proteomic analyses (liquid chromatography – mass spectrometry)

The primary sequence of peptides from IEC inhibitory subfractions were determined using liquid chromatography – mass spectrometry (LC-MS). Reconstituted subfractions were separated using reversed phase LC on an Ultimate 3000 chromatography system (Thermo Scientific, Sunnyvale, CA, USA). Peptides were loaded onto a C18 trap column (Thermo Scientific) and separated on a 150 mm PepMap C18 column (150 mm length, 75 μm i.d., 2 μm particle size, Thermo Scientific) heated to 65° C using a 30 minute gradient from 2 to 35% mobile phase B (95% acetonitrile, 0.2% formic acid) followed by column wash (at 85% mobile phase B) and re-equilibration at 2% mobile phase A (5% acetonitrile (*v/v*), 0.2% formic acid (*v/v*)) for a total run time of 60 min. The outlet of the column was coupled to a nano electrospray ionization source. All mass spectrometry measurements were performed on an Orbitrap Eclipse instrument (Thermo Scientific, San Jose, CA, USA). Broadband spectra were recorded at 120,000 resolving power (at *m/z* 200). Dynamic exclusion was enabled for 60 sec. A top-speed cycle (duration: 3 sec) was used to quadrupole-select precursor cations (allowed charge states: 2+ to 7+) based on decreasing signal intensity. Tandem mass spectrometry was performed by higher-energy collisional dissociation (HCD) at a normalized collision energy (NCE) of 30%, with spectra recorded in the Orbitrap at 7,500 resolving power (at *m/z* 200). Mass spectrometry RAW files were searched in Proteome Discoverer (Thermo Scientific) using MS Amanda and SEQUEST algorithms, as well as in MaxQuant, against a database derived from the AZ bark scorpion (*C. sculpturatus*) venom gland transcriptome provided by D.R. Rokyta (Florida State University). Transcriptome sequencing and assembly was performed using methods previously described [66,67]. A 10 ppm and 0.5 Da mass tolerance was applied to the match of intact (i.e., precursor) and fragment ions, respectively. A 1% false-discovery rate cutoff was applied to the database search results. Peptides that were identified by all three database searches were subsequently compared based on presence/absence in bioactive (inhibitory) and non-bioactive (non-inhibitory) subfractions.

### 5.8. Search of Peptides Analogs

The protein BLAST website (http://blast.ncbi.nlm.nih.gov) was used for to search for proteins with similar primary structure to the proteins (peptides) identified in this study. Proteins with high similarity percentages (at least 60%) were identified from the database and aligned with proteins from this study. The SWISS-MODEL (http://swissmodel.expasy.org/) was used to propose the 3D-structure of peptides.

### 5.9. Statistics

Data were analyzed and plotted using GraphPad Prism 9 (GraphPad Software, Inc., San Diego, CA, USA). Summarized whole cell current data reported as the mean ± SEM of the OtNav1.8 current density. Summarized data were compared by the Student’s (two-tailed) *t* test. P < 0.05 was considered significant.

## Supplementary Materials

The following are available online at www.mdpi.com/xxx/s1, **Table S1**: Thirty-five peptides were identified from 19 subfractions of F7 and F11. **Figure S1**: Fractions 1 – 6, 8 – 10, 14, 15 and 17 had no effect on OtNav1.8 Na^+^ current. **Figure S2**: Subfractions 7B, 7D, 7G, 7H, 7I, 7J, 7K, and 7L had no significant effect on OtNav1.8 Na^+^ current. **Figure S3**: Subfractions 11A, 11B, 11C, 11D, 11F, 11G, and 11H had no significant effect on OtNav1.8 Na^+^ current. **Figure S4**: Inhibitory effects of subfraction 7F on OtNav1.8 channels are voltage dependent. **Figure S5**: Inhibitory effects of subfraction 7M on OtNav1.8 channels are voltage dependent. **Figure S6**: Inhibitory effects of subfraction 7N on OtNav1.8 channels are voltage dependent. **Figure S7**: Primary structure of NaTx-4, identified from AZ bark scorpion venom that inhibit Nav1.8 Na^+^ current. **Figure S8**: Primary structure of NaTx-36, identified from AZ bark scorpion venom that inhibit Nav1.8 Na^+^ current. **Figure S9**: Primary structure of NaTx-13, identified from AZ bark scorpion venom that inhibit Nav1.8 Na^+^ current.

## Author Contributions

Conceptualization, T.M.A., Y.X. and A.H.R.; methodology and data collection, T.M.A., Y.X., L.F., J.K., K.D.L., H.G., A.H., D.R.R., M.J.W. and A.H.R.; software, T.M.A., Y.X., L.F., D.R.R., M.J.W. and A.H.R.; validation, T.M.A., Y.X., L.F., J.K., D.R.R., M.J.W. and A.H.R.; formal analysis, T.M.A., Y.X., L.F., J.K. and A.H.R.; resources, T.R.C., J.D.S., L.F. and A.H.R.; data curation, L.F., J.K., D.R.R., M.J.W., and A.H.R.; writing-original draft preparation, T.M.A. and A.H.R.; writing-review and editing, T.M.A, Y.X., J.K., H.G., A.H., K.D.L., M.J.W., D.R.R., J.D.S., T.R.C., L.F. and A.H.R.; visualization, T.M.A., L.F., J.K., and A.H.R.; supervision, A.H.R.; project administration, A.H.R.; funding acquisition, A.H.R. All authors have read and agreed to the published version of the manuscript.

## Acknowledgments

Thanks to Dr. Matthew P. Rowe for assistance collecting scorpions and extracting venom; to Dr. Bret Blum and the Santa Rita Experimental Station, University of Arizona for permission to collect specimens of C. *sculpturatus*; to Dr. Irina Vetter for comments on this manuscript; and to Dr. Ann West, Dr. Christina Bourne, and Dr. Robert Cichewicz for suggestions related to project methodology.

## References

1. Basbaum, A.I.; Bautista, D.M.; Scherrer, G.; Julius, D. Cellular and Molecular Mechanisms of Pain. Cell 2009, 139, 267–284, doi:10.1016/j.cell.2009.09.028.

2. Cavanaugh, D.J.; Lee, H.; Lo, L.; Shields, S.D.; Zylka, M.J.; Basbaum, A.I.; Anderson, D.J. Distinct subsets of unmyelinated primary sensory fibers mediate behavioral responses to noxious thermal and mechanical stimuli. Proceedings of the National Academy of Sciences 2009, 106, 9075–9080, doi:10.1073/pnas.0901507106.

3. Le Pichon, C.E.; Chesler, A.T. The functional and anatomical dissection of somatosensory subpopulations using mouse genetics. Frontiers in Neuroanatomy 2014, 8, 1–18.

4. Peirs, C.; Seal, R.P. Neural Circuits for Pain: Recent Advances and Current Views. Science 2016, 354, 578–583.

5. Renganathan, M.; Cummins, T.R.; Waxman, S.G. Contribution of Na^+^ 1.8 Sodium Channels to Action Potential Electrogenesis in DRG Neurons. Journal of Neurophysiology 2001, 86, 629–640.

6. Blair, N.T.; Bean, B.P. Roles of Tetrodotoxin (TTX)-Sensitive Na^+^ Current, TTX-Resistant Na^+^ Current, and Ca2+ Current in the Action Potentials of Nociceptive Sensory Neurons. The Journal of Neuroscience 2002, 22, 10277–10290.

7. Cummins, T.; Sheets, P.; Waxman, S. The roles of sodium channels in nociception: Implications for mechanisms of pain. Mol Interv 2007, 131, 243–257.

8. Dib-Hajj, S.D.; Cummins, T.R.; Black, J.A.; Waxman, S.G. Sodium channels in normal and pathological pain. Annual Review of Neuroscience 2010, 33, 325–347.

9. Barbosa, C.; Cummins, T.R. Chapter Eighteen - Unusual Voltage-Gated Sodium Currents as Targets for Pain. In Current Topics in Membranes, Robert, J.F., Sergei Yu, N., Eds. Academic Press: 2016; Vol. Volume 78, pp. 599–638.

10. Han, C.; Huang, J.; Waxman, S.G. Sodium channel Nav1.8: emerging links to human disease. Neurology 2016, 86, 473–483, doi:doi: 10.1212/WNL.0000000002333.

11. Abrahamsen, B.; Zhao, J.; Asante, C.O.; Cendan, C.M.; Marsh, S.; Martinez-Barbera, J.P.; Nassar, M.A.; Dickenson, A.H.; Wood, J.N. The Cell and Molecular Basis of Mechanical, Cold, and Inflammatory Pain. Science 2008, 321, 702–705, doi:10.1126/science.1156916.

12. Cummins, T.R.; Waxman, S.G. Downregulation of tetrodotoxin-resistant sodium currents and upregulation of a rapidly repriming tetrodotoxin-sensitive sodium current in small spinal sensory neurons after nerve injury. Journal of Neuroscience 1997, 17, 3503–3514.

13. Dib-Hajj, S.D.; Fjell, J.; Cummins, T.R.; Zheng, Z.; Fried, K.; LaMotte, R.; Waxman, S.G. Plasticity of sodium channel expression in DRG neurons in the chronic constriction injury model of neuropathic pain. Pain 1999, 83, 591–600.

14. Gold, M.S.; Reichling, D.B.; Shuster, M.J.; Levine, J.D. Hyperalgesic agents increase a tetrodotoxin-resistant Na^+^ current in nociceptors. Proceedings of the National Academy of Sciences 1996, 93, 1108–1112.

15. Okuse, K.; Chaplan, S.R.; McMahon, S.B.; Luo, Z.D.; Calcutt, N.A.; Scott, B.P.; Wood, J.N. Regulation of expression of the sensory neuron-specific sodium channel SNS in inflammatory and neuropathic pain. Molecular and Cellular Neuroscience 1997, 10, 196–207.

16. Tanaka, M.; Cummins, T.R.; Ishikawa, K.; Dib-Hajj, S.D.; Black, J.A.; Waxman, S.G. SNS Na^+^ channel expression increases in dorsal root ganglion neurons in the carrageenan inflammatory pain model. NeuroReport 1998, 9, 967–972.

17. Faber, C.G.; Lauria, G.; Merkies, I.S.; Cheng, X.; Han, C.; Ahn, H.S.; Waxman, S.G. Gain-of-function Nav1.8 mutations in painful neuropathy. Proceedings of the National Academy of Sciences 2012, 109, 19444–19449.

18. Han, C.; Vasylyev, D.; Macala, L.J.; Gerrits, M.M.; Hoeijmakers, J.G.; Bekelaar, K.J.; Waxman, S.G. The G1662S Nav1.8 mutation in small fibre neuropathy: impaired inactivation underlying DRG neuron hyperexcitability. Journal of Neurology, Neurosurgery, and Psychiatry 2014, 85, 499–505. 19.

19. Catterall, W.A. Cellular and molecular biology of voltage-gated sodium channels. Physiological Reviews 1992, 72, S15–S48.

20. Catterall, W.A. From ion currents to molecular mechanisms: the structure and function of voltage-gated sodium channels. Neuron 2000, 26, 13–25.

21. Ahern, C.A.; Payandeh, J.; Bosmans, F.; Chanda, B. The hitchhiker’s guide to the voltage-gated sodium channel galaxy. J Gen Physiol 2016, 147, 1–24, doi:10.1085/jgp.201511492.

22. Catterall, W.A. Structure and function of voltage-gated sodium channels at atomic resolution. Experimental Physiology 2014, 99, 1–26, doi:doi:10.1113/expphysiol.2013.071969.

23. Vilin, Y.Y.; Ruben, P.C. Slow inactivation in voltage-gated sodium channels. Cell Biochemistry and Biophysics 2001, 35, 171–190.

24. Green, B.R.; Olivera, B.M. Chapter Three - Venom Peptides From Cone Snails: Pharmacological Probes for Voltage-Gated Sodium Channels. In Current Topics in Membranes, Robert, J.F., Sergei Yu, N., Eds. Academic Press: 2016; Vol. Volume 78, pp. 65–86.

25. Maertens, C.; Cuypers, E.; Amininasab, M.; Jalali, A.; Vatanour, H.; Tytgat, J. Potent modulation of the voltage-gated sodium channel Nav1.7 by OD1, a toxin from the scorpion Odonthobuthus doriae Molecular Pharmacology 2006, 70, 405–414.

26. Osteen, J.D.; Herzig, V.; Gilchrist, J.; Emrick, J.J.; Zhang, C.; Wang, X.; Castro, J.; Garcia-Caraballo, S.; Grundy, L.; Rychkov, G.Y., et al. Selective spider toxins reveal a role for the Na^sub v^1.1 channel in mechanical pain. Nature 2016, 534, 494–499J, doi:http://dx.doi.org/10.1038/nature17976.

27. Rowe, A.H.; Xiao, Y.; Scales, J.; Linse, K.D.; Rowe, M.P.; Cummins, T.R.; Zakon, H.H. Isolation and characterization of CvIV4: a pain inducing alpha-scorpion toxin. PLoS One 2011, 6, e23520, doi:10.1371/journal.pone.0023520.

28. Vandendriessche, T.; Olamendi-Portugal, T.; Zamudio, F.Z.; Possani, L.D.; Tytgat, J. Isolation and characterization of two novel scorpoin toxins: The α-toxin-like CeII8, specific for Nav1.7 channels and the classical anti-mammalian CeII9, specific for Nav1.4 channels. Toxicon 2010, 56, 613–623.

29. Xiao, Y.; Bingham, J.-P.; Zhu, W.; Moczydlowski, E.; Liang, S.; Cummins, T.R. Tarantula Huwentoxin-IV Inhibits Neuronal Sodium Channels by Binding to Receptor Site 4 and Trapping the Domain II Voltage Sensor in the Closed Configuration. Journal of Biological Chemistry 2008, 283, 27300–27313, doi:10.1074/jbc.M708447200.

30. Xiao, Y.; Blumenthal, K.; Jackson, J.O.; Liang, S.; Cummins, T.R. The Tarantula Toxins ProTx-II and Huwentoxin-IV Differentially Interact with Human Nav1.7 Voltage Sensors to Inhibit Channel Activation and Inactivation. Molecular Pharmacology 2010, 78, 1124–1134, doi:10.1124/mol.110.066332.

31. Mohamed Abd El-Aziz, T.; Soares, A.G.; Stockand, J.D. Snake Venoms in Drug Discovery: Valuable Therapeutic Tools for Life Saving. Toxins 2019, 11, 564.

32. Bordon, K.d.C.F.; Cologna, C.T.; Fornari-Baldo, E.C.; Pinheiro-Júnior, E.L.; Cerni, F.A.; Amorim, F.G.; Anjolette, F.A.P.; Cordeiro, F.A.; Wiezel, G.A.; Cardoso, I.A., et al. From Animal Poisons and Venoms to Medicines: Achievements, Challenges and Perspectives in Drug Discovery. Frontiers in Pharmacology 2020, 11, doi:10.3389/fphar.2020.01132.

33. Pennington, M.W.; Czerwinski, A.; Norton, R.S. Peptide therapeutics from venom: Current status and potential. Bioorganic & medicinal chemistry 2018, 26, 2738–2758, doi:https://doi.org/10.1016/j.bmc.2017.09.029.

34. Lazarovici, P. Snake- and Spider-Venom-Derived Toxins as Lead Compounds for Drug Development. Methods Mol Biol 2020, 2068, 3–26, doi:10.1007/978-1-4939-9845-6_1.

35. McGivern, J.G. Ziconotide: a review of its pharmacology and use in the treatment of pain. Neuropsychiatr Dis Treat 2007, 3, 69–85, doi:10.2147/nedt.2007.3.1.69.

36. McDermott, A. News Feature: Venom back in vogue as a wellspring for drug candidates. Proceedings of the National Academy of Sciences 2020, 117, 10100, doi:10.1073/pnas.2004486117.

37. Flinspach, M.; Xu, Q.; Piekarz, A.D.; Fellows, R.; Hagan, R.; Gibbs, A.; Liu, Y.; Neff, R.A.; Freedman, J.; Eckert, W.A., et al. Insensitivity to pain induced by a potent selective closed-state Nav1.7 inhibitor. Scientific Reports 2017, 7, 39662, doi:10.1038/srep39662.

38. Shilong Yang, Y.X., Di Kang, Jie Liu, Yuan Li, Eivind A.B. Undheim, Julie K. Klint, Mingqiang Rong, Ren Lai, Glenn F. King. Discovery of a selective Nav1.7 inhibitor from centipede venom with analgesic efficacy exceeding morphine in rodent pain models. Proceedings of the National Academy of Sciences 2013, 110, 17534–17539.

39. Yin, K., Deuis, Jennifer .R., Dekan, Zoltan, Jin, Ai-Hua, Alewood, Paul, F., King, Glenn, F., Herzig, Volker, and Vetter, Irina. Addition of K22 Converts Spider Venom Peptide Pme2a from an Activator to an Inhibitor of Nav1.7. Biomedicines 2020, 8, 1–9, doi:10.3390/biomedicines8020037.

40. Selvaraj, U.M.; Zuurbier, K.R.; Whoolery, C.W.; Plautz, E.J.; Chambliss, K.L.; Kong, X.; Zhang, S.; Kim, S.H.; Katzenellenbogen, B.S.; Katzenellenbogen, J.A., et al. Selective Nonnuclear Estrogen Receptor Activation Decreases Stroke Severity and Promotes Functional Recovery in Female Mice. Endocrinology 2018, 159, 3848–3859, doi:10.1210/en.2018-00600.

41. Catterall, W.A.; Cestèle, S.; Yarov-Yarovoy, V.; Yu, F.H.; Konoki, K.; Scheuer, T. Voltage-gated ion channels and gating modifier toxins. Toxicon 2007, 49, 124–141, doi:http://dx.doi.org/10.1016/j.toxicon.2006.09.022.

42. Bosmans, F.; Swartz, K.J. Targeting voltage sensors in sodium channels with spider toxins. Trends in Pharmacological Sciences 2010, 31, 175–182, doi:10.1016/j.tips.2009.12.007.

43. Green, B.R.B. Venom Peptides From Cone Snails: Pharmacological Probes for Voltage-Gated Sodium Channels. Current topics in membranes 2016, 78, 65–86.

44. Zhang, J.Z.; Yarov-Yarovoy, V.; Scheuer, T.; Karbat, I.; Cohen, L.; Gordon, D.; Gurevitz, M.; Catterall, W.A. Structure-Function Map of the Receptor Site for beta-Scorpion Toxins in Domain II of Voltage-gated Sodium Channels. Journal of Biological Chemistry 2011, 286, 33641–33651, doi:10.1074/jbc.M111.282509.

45. Cestèle, S.; Qu, Y.; Rogers, J.C.; Rochat, H.; Scheuer, T.; Catterall, W.A. Voltage sensor-trapping: enhanced activation of sodium channels by beta-scorpion toxin bound to the S3-S4 loop in domain II. Neuron 1998, 21, 919–931, doi:10.1016/S0896-6273(00)80606-6.

46. Bosmans, F.; Tytgat, J. Voltage-gated sodium channel modulation by scorpion α-toxins. Toxicon 2007, 49, 142–158.

47. Clairfeuille, T.; Cloake, A.; Infield, D.T.; Llongueras, J.P.; Arthur, C.P.; Li, Z.R.; Jian, Y.W.; Martin-Eauclaire, M.F.; Bougis, P.E.; Ciferri, C., et al. Structural basis of alpha-scorpion toxin action on Na-v channels. Science 2019, 363, 1302–+, doi:10.1126/science.aav8573.

48. Gilchrist, J.; Bosmans, F. Using voltage-sensor toxins and their molecular targets to investigate NaV1.8 gating. The Journal of Physiology 2018, 596, 1863–1872, doi:10.1113/jp275102.

49. Rowe, A.H.; Xiao, Y.; Rowe, M.P.; Cummins, T.R.; Zakon, H.H. Voltage-gated sodium channel in grasshopper mice defends against bark scorpion toxin. Science 2013, 342, 441–446, doi:10.1126/science.1236451.

50. Couraud, F.; Jover, E. Mechanism of Action of Scorpion Toxins. In Handbook of Natural Toxins, Tu, A.T., Ed. Marcel Dekker, Inc.: New York, N.Y., 1984; Vol. 2, pp. 659–678.

51. Corona, M.; Valdez-Cruz, N.A.; Merino, E.; Zurita, M.; Possani, L.D. Genes and peptides from the scorpion Centruroides sculpturatus Ewing, that recognize Na^+^-channels. Toxicon 2001, 39, 1893–1898.

52. Possani, L.D.; Becerril, B.; Delepierre, M.; Tytgat, J. Scorpion toxins specific for Na^+^-channels. European Journal of Biochemistry 1999, 264, 287–300.

53. Possani, L.D.; Merino, E.; Corona, M.; Bolivar, F.; Becerril, B. Peptides and genes coding for scorpion toxins that affect ion-channels. Biochimie 2000, 82, 861–868.

54. Rodriguez de la Vega, R.C.; Possani, L.D. Current views on scorpion toxins specific for K^+^ - channels. Toxicon 2004, 43, 865–875.

55. Rodriguez de la Vega, R.; Possani, L.D. Overview of scorpion toxins specific for Na^+^ channels and related peptides: biodiversity, structure-function relationships and evolution. Toxicon 2005, 46, 831–844.

56. Simard, J.M.; Meves, H.; Watt, D.D. Neurotoxins in venom from the North American scorpion, Centruroides sculpturatus Ewing. In Natural Toxins: Toxicolgy, Chemistry and Safety, Keeler, R.F., Mandava, N.B., Tu, A.T., Eds. Alaken, Inc.: Fort Collins, CO, 1992; Vol. 1, pp. 236–263.

57. Rowe, A.H.; Rowe, M.P. Risk assessment by grasshopper mice (Onychomys spp.) feeding on neurotoxic prey (Centruroides spp.). Animal Behaviour 2006, 71, 725–734.

58. Rowe, A.H.; Rowe, M.P. Physiological resistance of grasshopper mice (Onychomys spp.) to Arizona bark scorpion (Centruroides exilicauda) venom. Toxicon 2008, 52, 597–605.

59. Zhang, F.; Zhang, C.; Xu, X.; Zhang, Y.; Gong, X.; Yang, Z.; Zhang, H.; Tang, D.; Liang, S.; Liu, Z. Naja atra venom peptide reduces pain by selectively blocking the voltage-gated sodium channel Nav1.8. J Biol Chem 2019, 294, 7324–7334, doi:10.1074/jbc.RA118.007370.

60. Valdez-Cruz, N.A.; Dávila, S.; Licea, A.; Corona, M.; Zamudio, F.Z.; García-Valdes, J.; Boyer, L.; Possani, L.D. Biochemical, genetic and physiological characterization of venom components from two species of scorpions: Centruroides exilicauda Wood and Centruroides sculpturatus Ewing. Biochimie 2004, 86, 387–396, doi:https://doi.org/10.1016/j.biochi.2004.05.005.

61. Yaksh, T.L.; Woller, S.A.; Ramachandran, R.; Sorkin, L.S. The search for novel analgesics: targets and mechanisms. F1000Prime Reports 2015, 7, 1–27, doi:doi:10.12703/P7-56.

62. Zhorov, B.S.; Tikhonov, D.B. Chapter Five - Computational Structural Pharmacology and Toxicology of Voltage-Gated Sodium Channels. In Current Topics in Membranes, Robert, J.F., Sergei Yu, N., Eds. Academic Press: 2016; Vol. Volume 78, pp. 117–144.

63. French, R.J.; Yoshikami, D.; Sheets, M.F.; Olivera, B.M. The Tetrodotoxin Receptor of Voltage-Gated Sodium Channels—Perspectives from Interactions with μ-Conotoxins. Marine Drugs 2010, 8, 2153–2161, doi:10.3390/md8072153.

64. Rogers, J.C.; Qu, Y.; Tanada, T.N.; Scheuer, T.; Catterall, W.A. Molecular determinants of high affinity binding of α-scorpion toxin and sea anemone toxin in the S3-S4 extracellular loop in domain IV of the Na^+^ channel α subunit. The Journal of Biological Chemistry 1996, 271, 15950–15962.

65. Flinspach, M.; Xu, Q.; Piekarz, A.D.; Fellows, R.; Hagan, R.; Gibbs, A.; Liu, Y.; Neff, R.A.; Freedman, J.; Eckert, W.A., et al. Insensitivity to pain induced by a potent selective closed-state Nav1.7 inhibitor. Sci Rep 2017, 7, 39662, doi:10.1038/srep39662.

66. Rokyta, D.R., and Ward, Micaiah J. Venom-gland transcriptomics and venom proteomics of the black-back scorpion (Hadrurus spadix) reveal detectability challenges and an unexplored realm of animal toxin diversity. Toxicon 2017, 128, 23–37.

67. Ward, M.J., Ellsworth, Schyler A., and Rokyta, Darin R. Venom-gland transcriptomics and venom proteomics of the Hentz striped scorpion (Centruroides hentzi; Buthidae) reveal high toxin diversity in a harmless member of a lethal family. Toxicon 2018, 142, 14–29.

