## Supplementary material for "Identification and characterization of novel proteins from Arizona bark scorpion venom that inhibit Nav1.8, a voltage-gated sodium channel regulator of pain signaling": Supplementary Material.docx

**Table S1.** Thirty five peptides were identified from 19 subfractions of F7 and F11.

| Fraction | Subfraction | Total Number of Toxins Identified in Fraction | Number of Toxins Unique to Fraction | Identified Toxins |
| --- | --- | --- | --- | --- |
| 7 | A* | 0 | 0 | Ψ |
|  | B | 0 | 0 | Ψ |
|  | C* | 2 | 1 | VP-2, NaTx-3 |
|  | D | 0 | 0 | Ψ |
|  | E* | 0 | 0 | Ψ |
|  | G | 4 | 2 | KTx-14, aNaTx-7, KTx-13, aNaTx-3 |
|  | H | 6 | 1 | NaTx-18, aKTx-2, VP-16, NaTx-7, NaTx-12, KTx-14 |
|  | J | 6 | 2 | SP-3, KTx-16, NaTx-12, NaTx-18, NaTx-7, aKTx-2 |
|  | K | 3 | 1 | aKTx-2, NaTx-18, bNaTx-13 |
| 11 | A | 0 | 0 | Ψ |
|  | B | 0 | 0 | Ψ |
|  | C | 0 | 0 | Ψ |
|  | D | 5 | 2 | VP-16, VP-2, NaTx-3, bNaTx-18, bNaTx-20 |
|  | E* | 6 | 4 | NaTx-4, NaTx-13, bNaTx-7, NaTx-22,  NaTx-36, VP-16 |
|  | F | 0 | 0 | Ψ |
|  | G | 0 | 0 | Ψ |
|  | H | 2 | 0 | bNaTx-15, bNaTx-7 |
|  | I* | 1 | 0 | bNaTx-15 |
|  | J* | 0 | 0 | Ψ |
| Total | | 35 | 13 | Number of Unique Toxins Identified: 22 |
| Toxins exclusive to bioactive (OtNav1.8-inhibiting) subfractions: NaTx-4, NaTx-13, NaTx-22, NaTx-36 | | | | |
| * Indicates bioactive subfraction | | | | |
| Ψ Indicates that not enough protein was in the sample to identify peptide sequences | | | | |

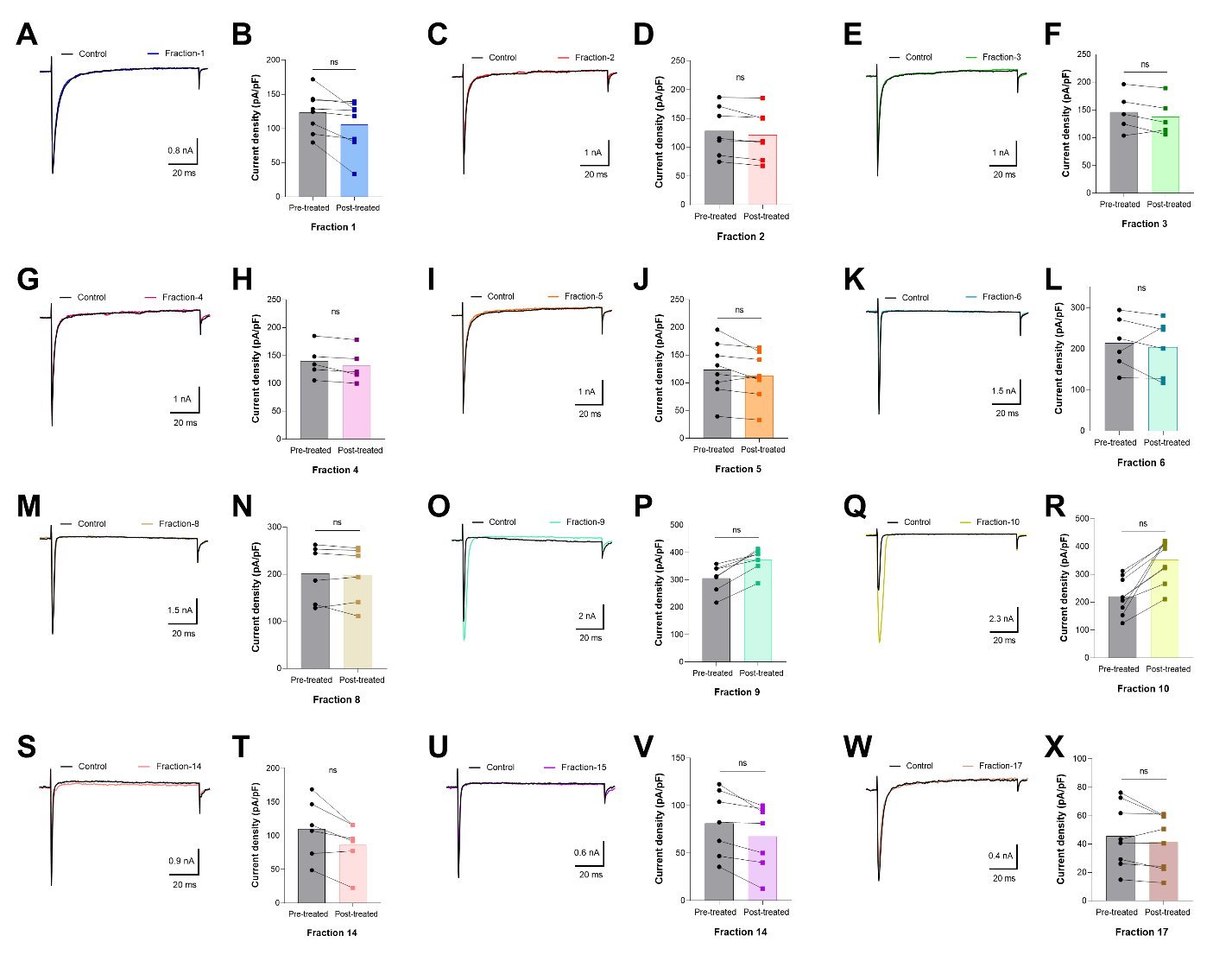

**Figure S1.** Fractions 1 – 6, 8 – 10, 14, 15 and 17 had no effect on OtNav1.8 Na+ current. (**A, C, E, G, I, K, M, O, Q, S, U,** and **W**) Representative OtNav1.8 currents before (black traces) and after application of fractions 1 (blue trace), 2 (red trace), 3 (green trace), 4 (violet trace), 5 (orange trace), 6 (turquoise trace), 8 (light brown trace), 9 (light green trace), 10 (light cumin color trace), 14 (light pink trace), 15 (light violet trace), and 17 (brown trace). The currents were elicited by a 100-msec depolarization to +20 mV from a holding potential of -80 mV before and after application of fractions (0.1 – 0.3 µg/mL) 1 – 6, 8 – 10, 14, 15 and 17. (**B, D, F, H, J, L, N, P, R, T, V,** and **X**) Summary graphs of Na^+^ currents quantified in whole-cell voltage clamped ND7/23 cells transfected with OtNav1.8 before (black circles, gray bars) and after application of fractions 1 (blue squars, light blue bar), 2 (red squars, light red bar), 3 (green squars, light green bar), 4 (violet squars, light violet bar), 5 (orange squars, light orange bar), 6 (turquoise squars, light urquoise bar), 8 (brown squars, light brown bar), 9 (light turquoise squars, light turquoise bar), 10 (cumin squars, light cumin bar), 14 (pink squars, light pink bar), 15 (violet squars, light violet bar), and 17 (brown squars, light red bar). Summary data from experiments (n = 5 – 9 cells) identical to those shown in **A, C, E, G, I, K, M, O, Q, S, U,** and **W**. * P < 0.05 vs. before application of fractions 1 – 6, 8 – 10, 14, 15, and 17.

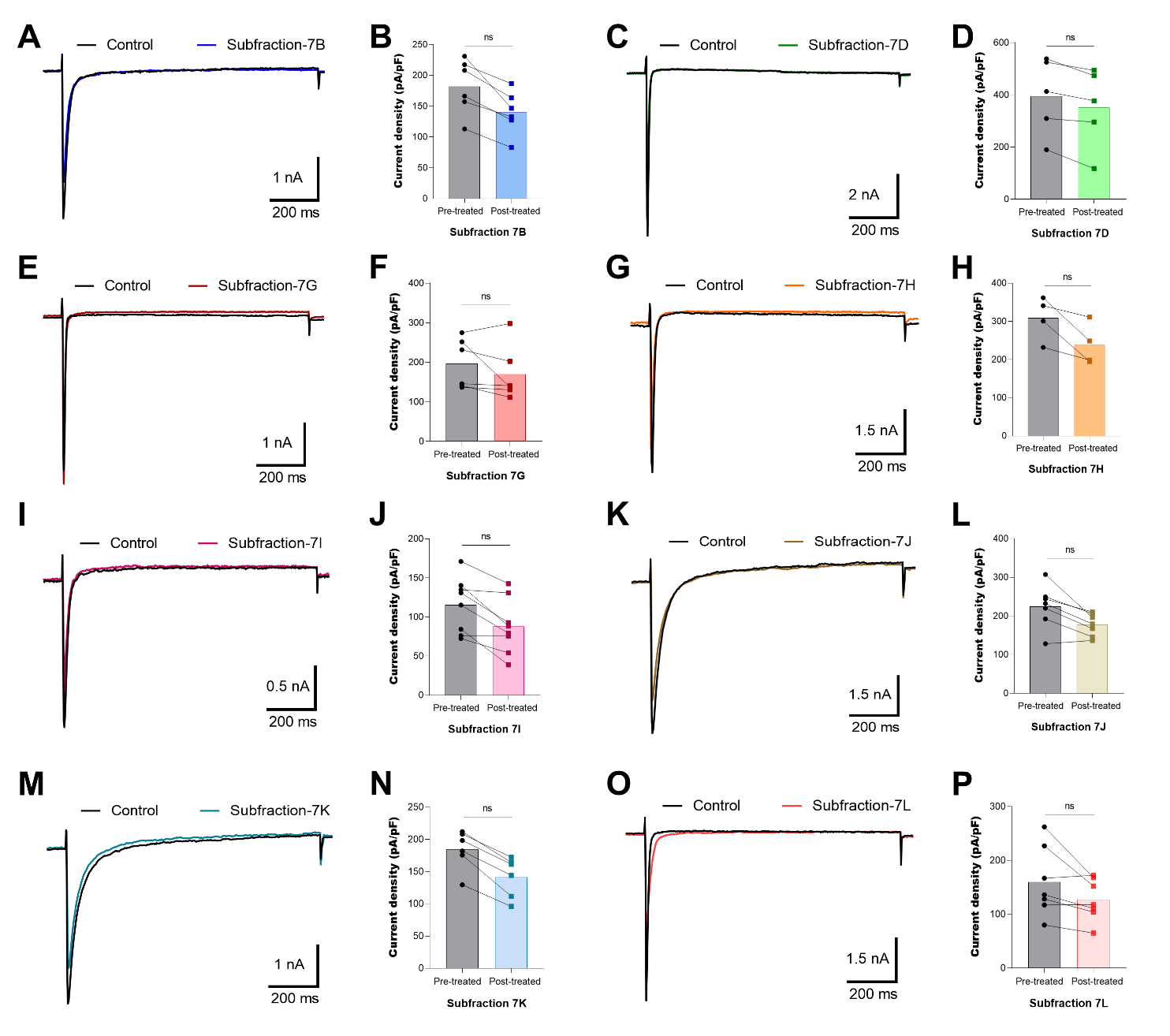

**Figure S2.** Subfractions 7B, 7D, 7G, 7H, 7I, 7J, 7K, and 7L had no significant effect on OtNav1.8 Na+ current. (**A, C, E, G, I, K, M,** and **O**) Representative OtNav1.8 currents before (black traces) and after application of subfractions 7B (blue trace), 7D (green trace), 7G (drak red trace), 7H (orange trace), 7I (violet trace), 7J (cumin color trace), 7K (turquoise trace), and 7L (red trace). The currents were elicited by a 100-msec depolarization to +20 mV from a holding potential of -80 mV before and after application of subfractions (0.1 – 0.3 µg/mL) 7B, 7D, 7G, 7H, 7I, 7J, 7K, and 7L. (**B, D, F, H, J, L, N,** and **P**) Summary graphs of Na^+^ currents quantified in whole-cell voltage clamped ND7/23 cells transfected with OtNav1.8 before (black circles, gray bars) and after application of subfractions 7B (blue squars, light blue bar), 7D (green squars, light green bar), 7G (drak red squars, red bar), 7H (orange squars, light orange bar), 7J (violet squars, light violet bar), 7L (cumin squars, light cumin bar), 7N (turquoise squars, light urquoise bar), and 7 P (red squars, light red bar). Summary data from experiments (n = 5 – 9 cells) identical to those shown in **A, C, E, G, I, K, M,** and **O**. * P < 0.05 vs. before application of subfractions 7B, 7D, 7G, 7H, 7I, 7J, 7K, and 7L.

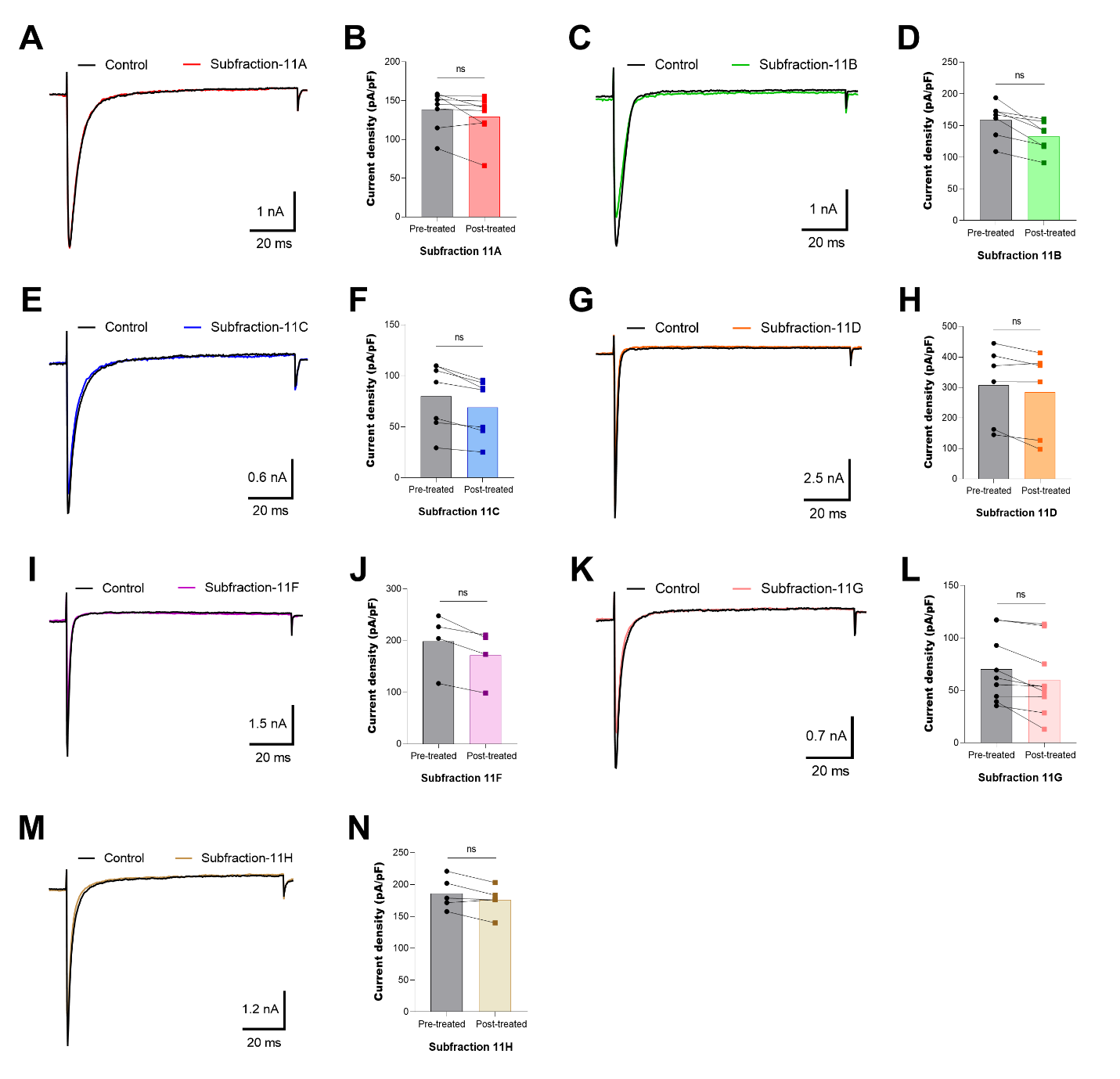

**Figure S3.** Subfractions 11A, 11B, 11C, 11D, 11F, 11G, and 11H had no significant effect on OtNav1.8 Na+ current. (**A, C, E, G, I, K,** and **M**) Representative OtNav1.8 currents before (black traces) and after application of subfractions 11A (red trace), 11B (green trace), 11C (blue trace), 11D (orange trace), 11F (violet trace), 11G (pink trace), and 11H (cumin color trace). The currents were elicited by a 100-msec depolarization to +20 mV from a holding potential of -80 mV before and after application of subfractions (0.1 – 0.3 µg/mL) 11A, 11B, 11C, 11D, 11F, 11G, and 11H. (**B, D, F, H, J, L,** and **N**) Summary graphs of Na^+^ currents quantified in whole-cell voltage clamped ND7/23 cells transfected with OtNav1.8 before (black circles, gray bars) and after application of 0.1-0.3 µg/mL of subfractions 11A (red squars, light red bar), 11B (green squars, light green bar), 11C (blue squars, light blue bar), 11D (orange squars, light orange bar), 11F (violet squars, light violet bar), 11G (pink squars, light pink bar), and 11H (cumin squars, light cumin bar). Summary data from experiments (n = 5 – 9 cells) identical to those shown in **A, C, E, G, I, K,** and **M**. * P < 0.05 vs. before application of subfractions 11A, 11B, 11C, 11D, 11F, 11G, and 11H.

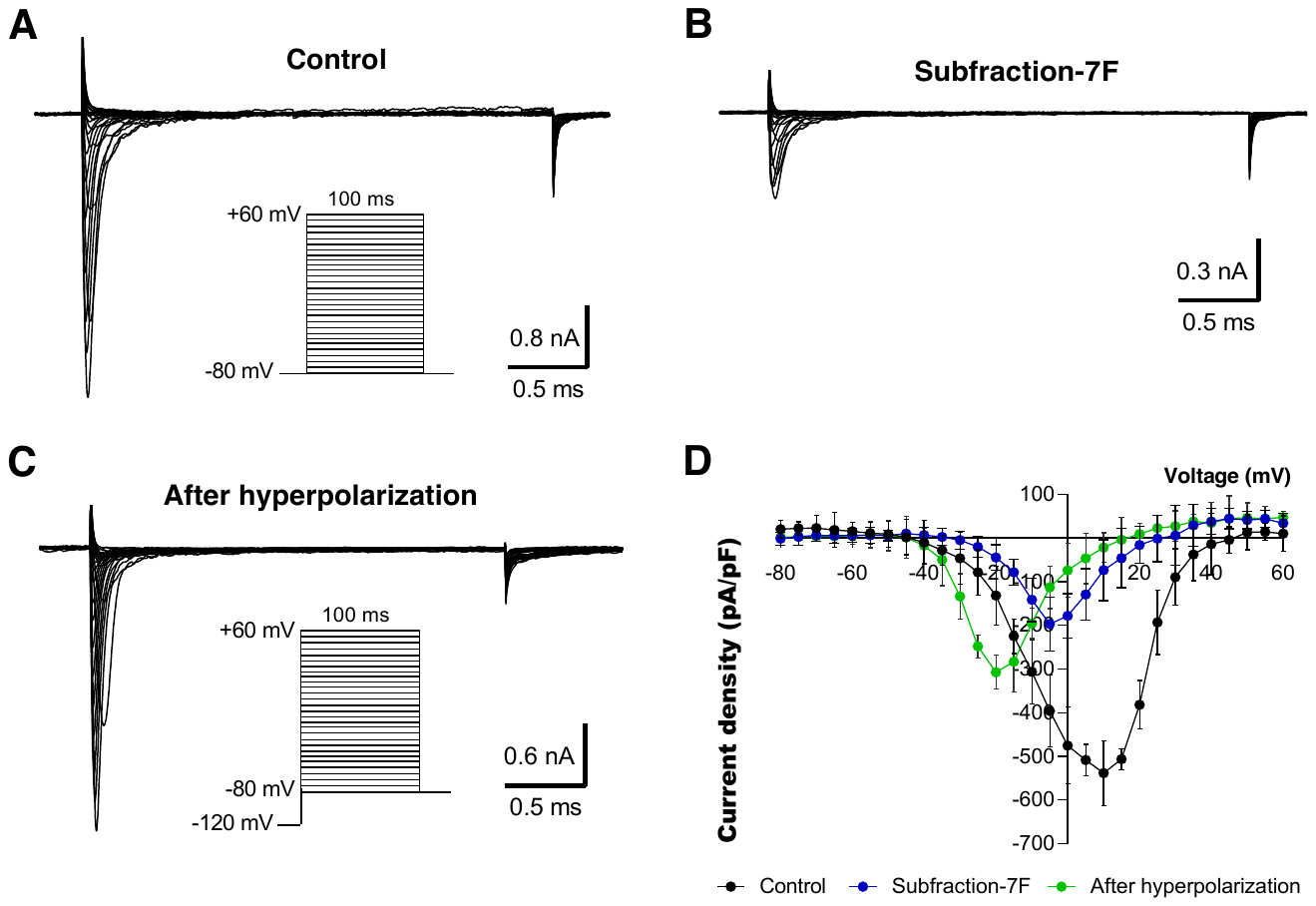

**Figure S4.** Inhibitory effects of subfraction 7F on OtNav1.8 channels are voltage dependent. (**A** and **B**) Representative activation current traces of OtNav1.8 channel expressed in ND7/23 cells in absence (control, **A**) in presence (**B**) of the subfraction 7F (0.1 – 0.3 µg/mL). The protocol is shown in inset. Total currents were elicited by a family of 100 ms depolarization from -80 mV to +60 mV in 5 mV increments. (**C**) Representative activation current traces of OtNav1.8 in presence of the subfraction 7F after hyperpolarizing the cell membranes (the protocol is shown in inset). The current-voltage curve was generated by voltage-clamp protocols consisting of holding potential of -120 mV for 30 sec followed a family of 100 ms depolarization from -80 mV to +60 mV in 5 mV increments. (**D**) I-V curves of the total Na^+^ currents in the absence (control, black circles) and presence of the subfraction 7F (blue circles) and after hyperpolarization (green circles) at a holding potential of -120 mV for 30 sec followed by a family of 100 ms depolarization from -80 mV to +60 mV in 5 mV increments. The inhibitory effects of subfraction 7F on Na^+^ current were reduced when ND7/23 cells (expressing OtNav1.8) were hyperpolarized to a holding potential of -120 mV compared to -80 mV. Summary data from experiments (n = 3 – 5 cells) identical to those shown in **A, B,** and **C**.

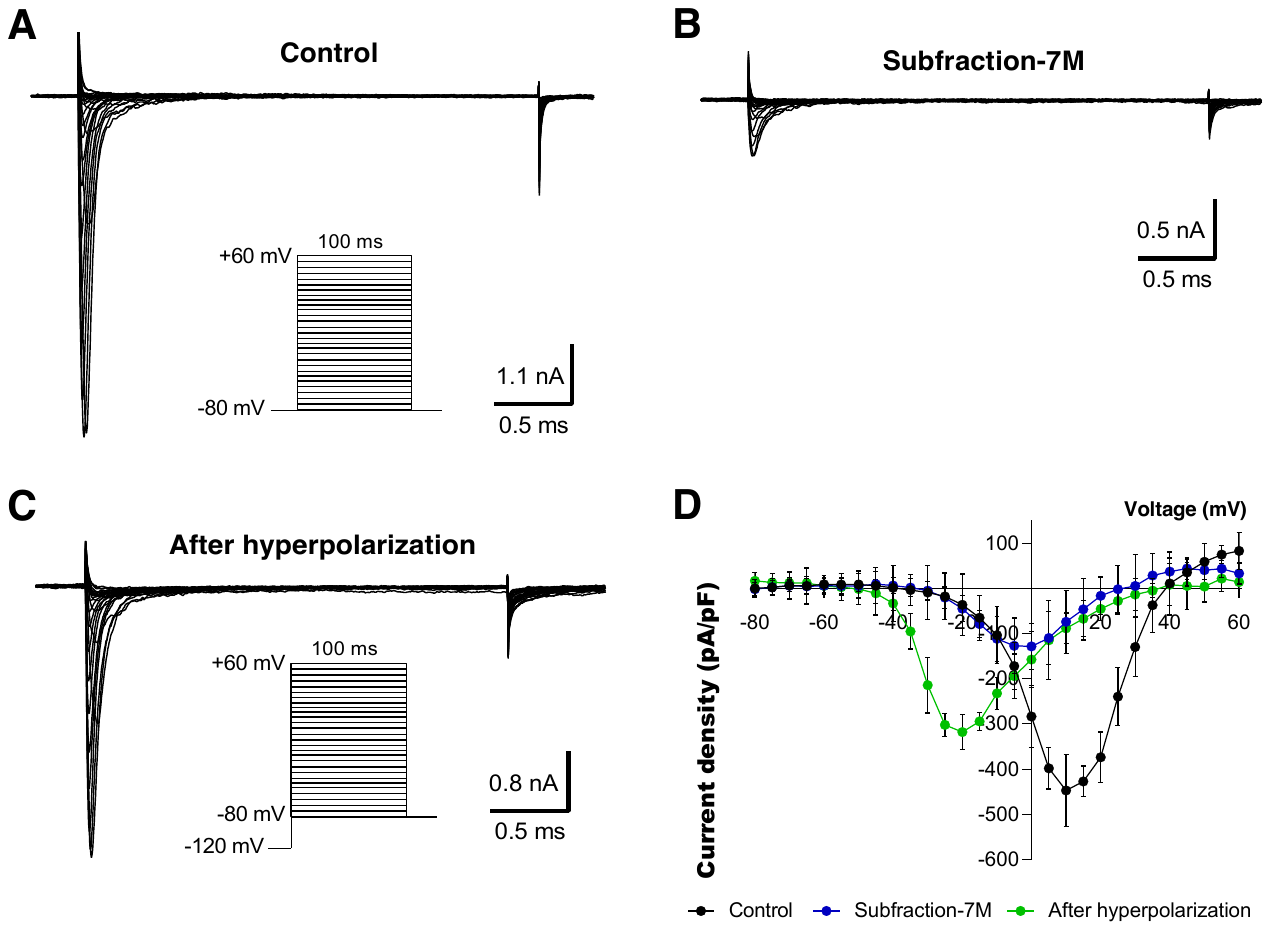

**Figure S5.** Inhibitory effects of subfraction 7M on OtNav1.8 channels are voltage dependent. (**A** and **B**) Representative activation current traces of OtNav1.8 channel expressed in ND7/23 cells in absence (control, **A**) in presence (**B**) of the subfraction 7M (0.1 – 0.3 µg/mL). The protocol is shown in inset. Total currents were elicited by a family of 100 ms depolarization from -80 mV to +60 mV in 5 mV increments. (**C**) Representative activation current traces of OtNav1.8 in presence of the subfraction 7M after hyperpolarizing the cell membranes (the protocol is shown in inset). The current-voltage curve was generated by voltage-clamp protocols consisting of holding potential of -120 mV for 30 sec followed a family of 100 ms depolarization from -80 mV to +60 mV in 5 mV increments. (**D**) I-V curves of the total Na^+^ currents in the absence (control, black circles) and presence of the subfraction 7M (blue circles) and after hyperpolarization (green circles) at a holding potential of -120 mV for 30 sec followed by a family of 100 ms depolarization from -80 mV to +60 mV in 5 mV increments. The inhibitory effects of subfraction 7M on Na^+^ current were reduced when ND7/23 cells (expressing OtNav1.8) were hyperpolarized to a holding potential of -120 mV compared to -80 mV. Summary data from experiments (n = 3 – 5 cells) identical to those shown in **A, B,** and **C**.

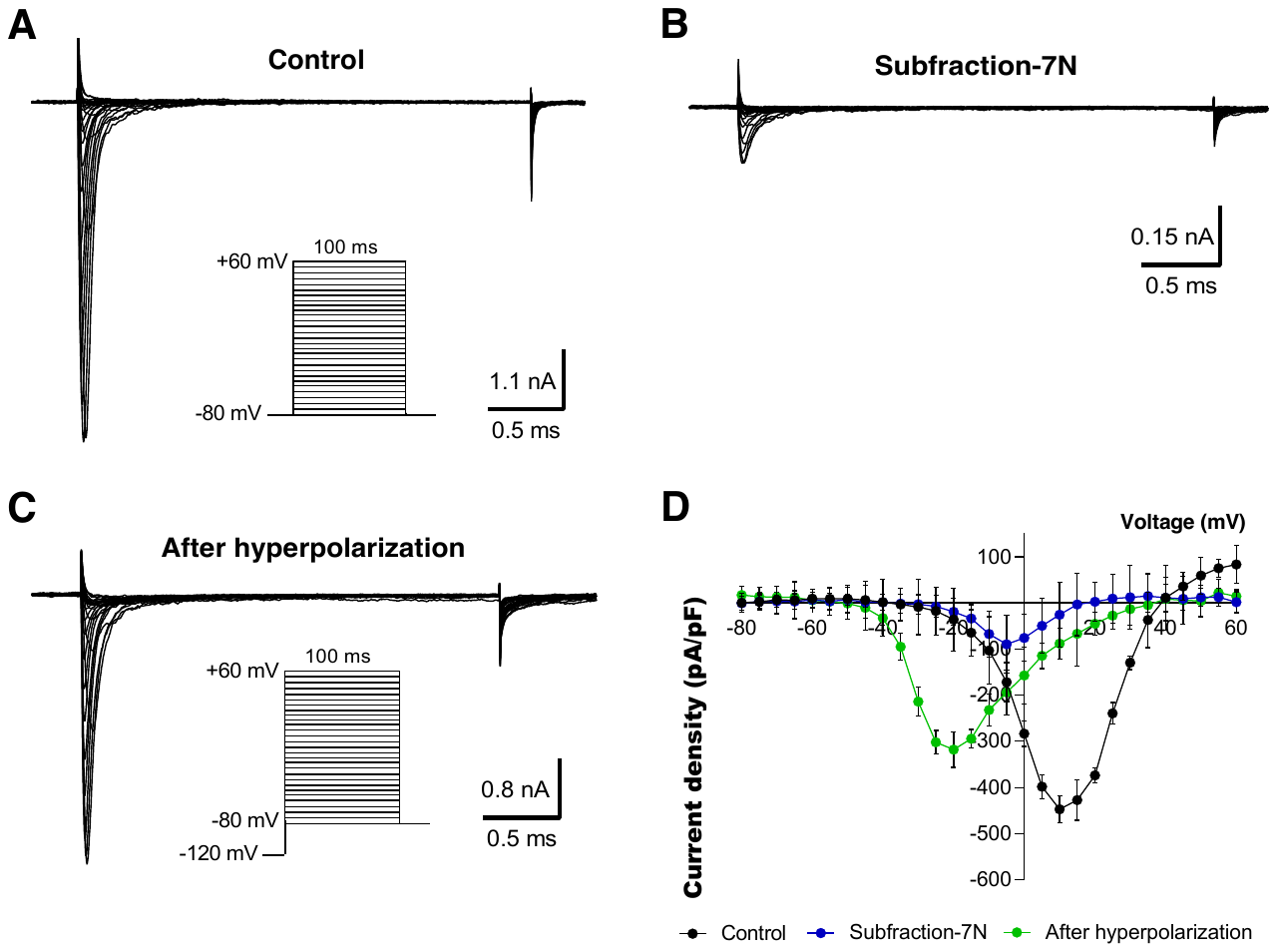

**Figure S6.** Inhibitory effects of subfraction 7N on OtNav1.8 channels are voltage dependent. (**A** and **B**) Representative activation current traces of OtNav1.8 channel expressed in ND7/23 cells in absence (control, **A**) in presence (**B**) of the subfraction 7N (0.1 – 0.3 µg/mL). The protocol is shown in inset. Total currents were elicited by a family of 100 ms depolarization from -80 mV to +60 mV in 5 mV increments. (**C**) Representative activation current traces of OtNav1.8 in presence of the subfraction 7N after hyperpolarizing the cell membranes (the protocol is shown in inset). The current-voltage curve was generated by voltage-clamp protocols consisting of holding potential of -120 mV for 30 sec followed a family of 100 ms depolarization from -80 mV to +60 mV in 5 mV increments. (**D**) I-V curves of the total Na^+^ currents in the absence (control, black circles) and presence of the subfraction 7N (blue circles) and after hyperpolarization (green circles) at a holding potential of -120 mV for 30 sec followed by a family of 100 ms depolarization from -80 mV to +60 mV in 5 mV increments. The inhibitory effects of subfraction 7N on Na^+^ current were reduced when ND7/23 cells (expressing OtNav1.8) were hyperpolarized to a holding potential of -120 mV compared to -80 mV. Summary data from experiments (n = 3 – 5 cells) identical to those shown in **A, B,** and **C**.

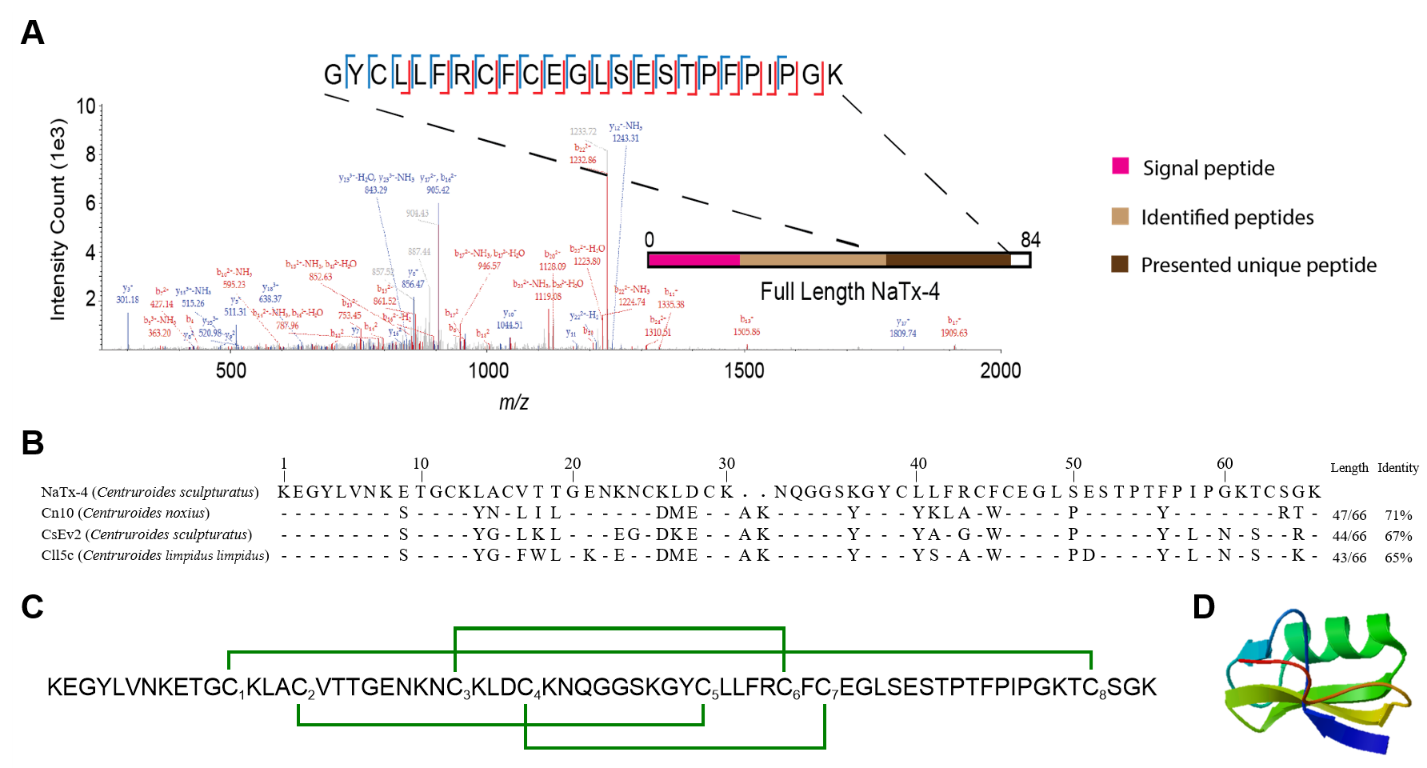

**Figure S7.** Primary structure of NaTx-4, identified from AZ bark scorpion venom that inhibit Nav1.8 Na+ current. (**A**) Tandem mass spectrometry-based identification of a unique peptide from NaTx-4. Experimental fragment ions matched against in silico-generated fragment ions (using a 0.5 Da mass tolerance) are indicated in red (*b*-ions, containing the N-terminus) or blue (*y*-ions, containing the C-terminus). Identified cleavage sites are recapitulated in the inset showing the exact sequence of the peptide. The position of the identified tryptic peptide within the whole sequence of NaTx-4 is indicated in brown on the bar representing the full sequence of the toxin protein. This tryptic peptide has a sequence unique to this toxin, and the intact mass of the peptide was matched with a 5.3 ppm deviation over the theoretical value. (**B**) Sequence alignment of NaTx-4 with homology toxins from NCBI BLAST. UniProt Knowledgebase (UniProtKB)/Swiss-Prot accession codes for these retrieved sequences are Q94435.2 (Cn10), P45663.1 (CsEv2), and Q7YT61.1 (CII5c). Hyphen-minus represents identical amino acid residues, and dots indicate the lack of residue at the position. The toxin lengths and percentages of sequence identities are given on the right. (**C**) Disulfide bridge arrangement (in green) of NaTx-4 proposed by homology with CsEv2. Cysteines were numbered by their order of appearance in the sequence. (**D**) SWISS-MODEL (http://swissmodel.expasy.org/) proposed the 3D-structure of NaTx-4.

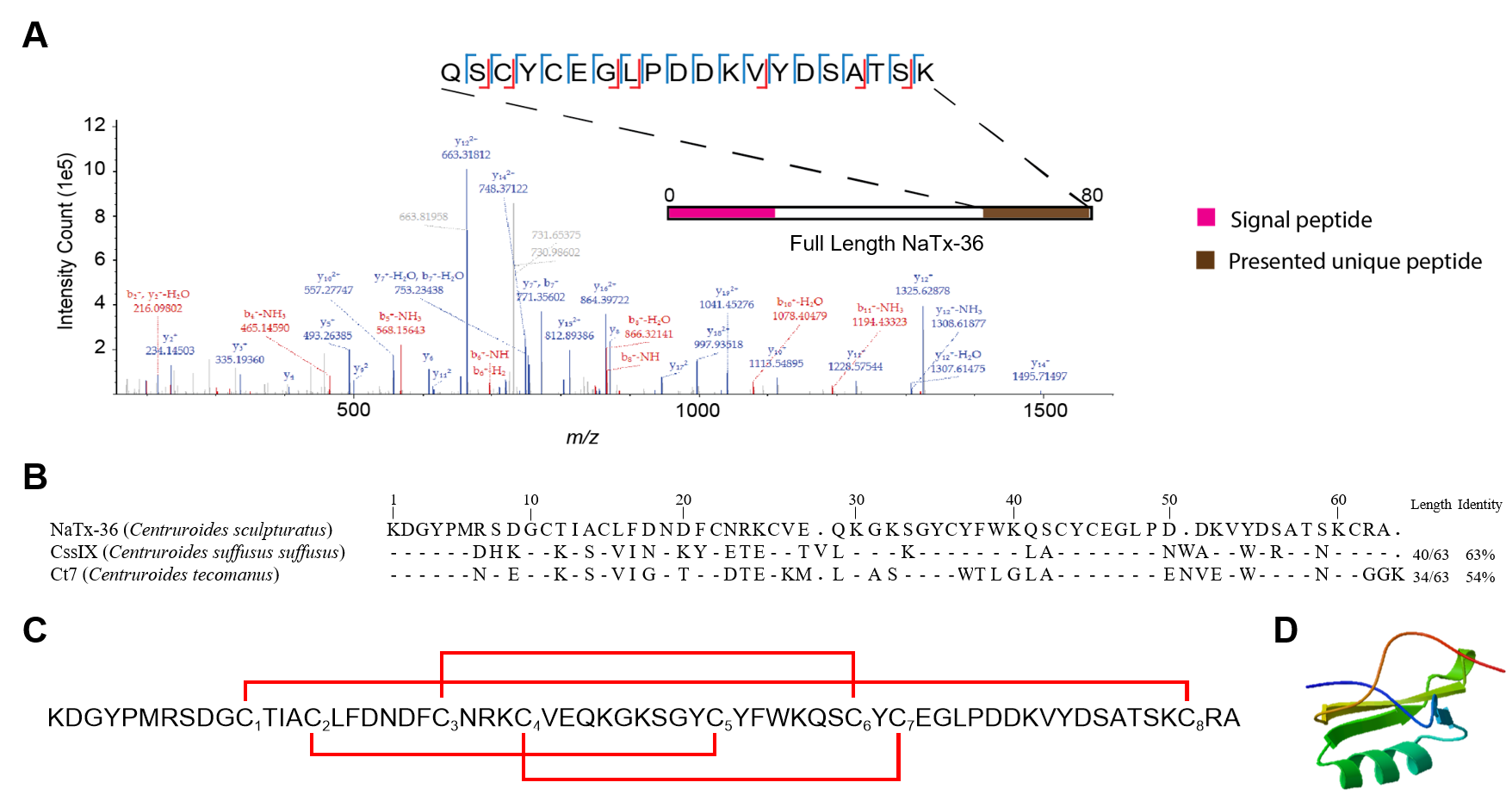

**Figure S8.** Primary structure of NaTx-36, identified from AZ bark scorpion venom that inhibit Nav1.8 Na+ current. (**A**) Tandem mass spectrometry-based identification of a unique peptide from NaTx-36. Experimental fragment ions matched against in silico-generated fragment ions (using a 0.5 Da mass tolerance) are indicated in red (*b*-ions, containing the N-terminus) or blue (*y*-ions, containing the C-terminus). Identified cleavage sites are recapitulated in the inset showing the exact sequence of the peptide. The position of the identified tryptic peptide within the whole sequence of NaTx-36 is indicated in brown on the bar representing the full sequence of the toxin protein. This tryptic peptide has a sequence unique to this toxin, and the intact mass of the peptide was matched with a 5.3 ppm deviation over the theoretical value. (**B**) Sequence alignment of NaTx-36 with homology toxins from NCBI BLAST. UniProt Knowledgebase (UniProtKB)/Swiss-Prot accession codes for these retrieved sequences are F1CGT6.1 (CssIX) and P0DUH9.1 (Ct7). Hyphen-minus represents identical amino acid residues, and dots indicate the lack of residue at the position. The toxin lengths and percentages of sequence identities are given on the right. (**C**) Disulfide bridge arrangement (in red) of NaTx-36 proposed by homology with CssIX. Cysteines were numbered by their order of appearance in the sequence. (**D**) SWISS-MODEL (http://swissmodel.expasy.org/) proposed the 3D-structure of NaTx-36.

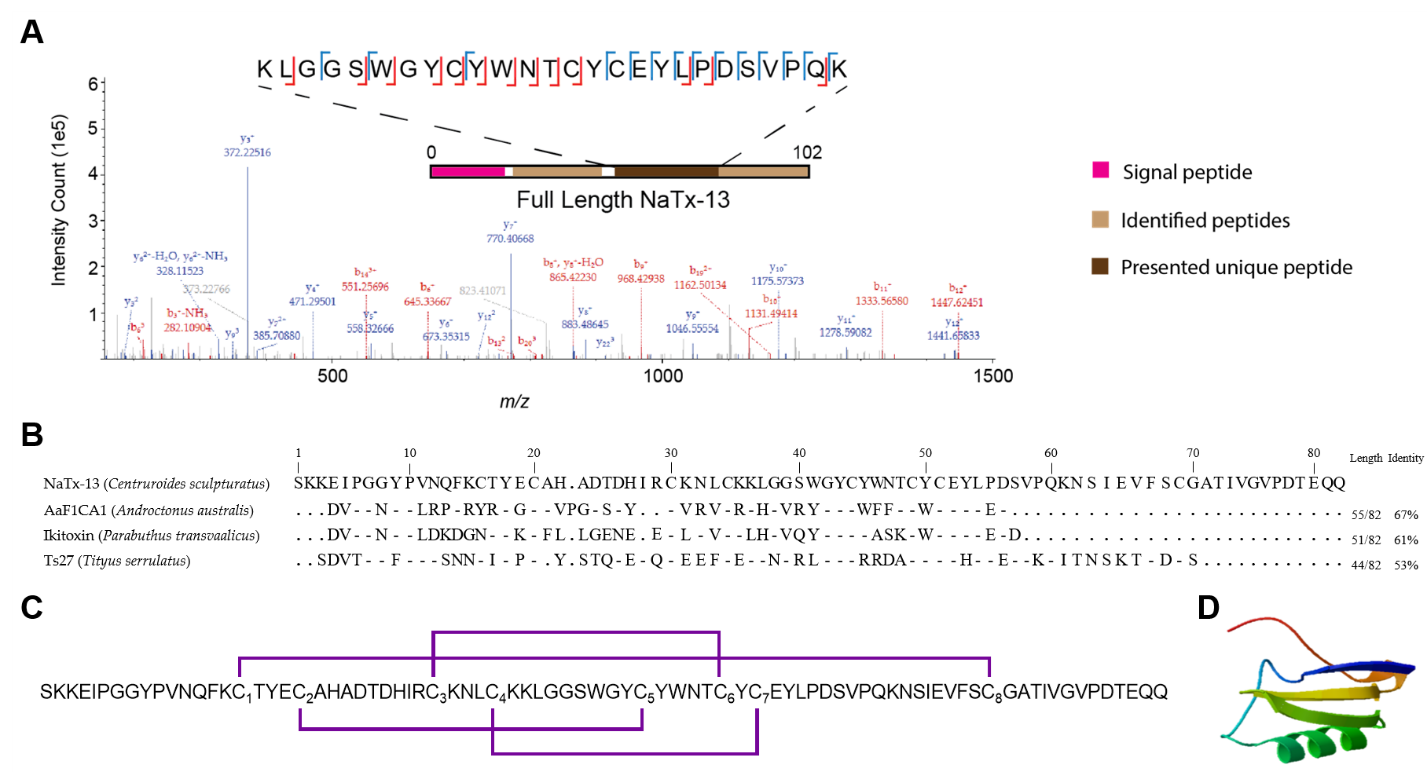

**Figure S9.** Primary structure of NaTx-13, identified from AZ bark scorpion venom that inhibit Nav1.8 Na+ current. (**A**) Tandem mass spectrometry-based identification of a unique peptide from NaTx-13. Experimental fragment ions matched against in silico-generated fragment ions (using a 0.5 Da mass tolerance) are indicated in red (*b*-ions, containing the N-terminus) or blue (*y*-ions, containing the C-terminus). Identified cleavage sites are recapitulated in the inset showing the exact sequence of the peptide. The position of the identified tryptic peptide within the whole sequence of NaTx-13 is indicated in brown on the bar representing the full sequence of the toxin protein. This tryptic peptide has a sequence unique to this toxin, and the intact mass of the peptide was matched with a 5.3 ppm deviation over the theoretical value. (**B**) Sequence alignment of NaTx-13 with homology toxins from NCBI BLAST. UniProt Knowledgebase (UniProtKB)/Swiss-Prot accession codes for these retrieved sequences are Q4LCT3.1 (AaF1CA1), P0C1B8.1 (Ikitoxin), and QPD99023.1 (Ts27). Hyphen-minus represents identical amino acid residues, and dots indicate the lack of residue at the position. The toxin lengths and percentages of sequence identities are given on the right. (**C**) Disulfide bridge arrangement (in violet) of NaTx-13 proposed by homology with AaF1CA1. Cysteines were numbered by their order of appearance in the sequence. (**D**) SWISS-MODEL (http://swissmodel.expasy.org/) proposed the 3D-structure of NaTx-13.
